# Structures of the Foamy virus fusion protein reveal an unexpected link with the F protein of paramyxo- and pneumoviruses

**DOI:** 10.1101/2024.02.09.579661

**Authors:** Ignacio Fernández, François Bontems, Delphine Brun, Youna Coquin, Casper A. Goverde, Bruno E. Coreilla, Antoine Gessain, Florence Buseyne, Felix A. Rey, Marija Backovic

## Abstract

Foamy viruses (FVs) constitute a subfamily of retroviruses. Their envelope glycoprotein (Env) drives the merger of viral and cellular membranes during entry into cells. The only available structures of retroviral Envs are those from human and simian immunodeficiency viruses from the subfamily of orthoretroviruses, which are only distantly related to the FVs. We report here the cryo-EM structures of the FV Env ectodomain in the pre- and post-fusion states, which demonstrate structural similarity with the fusion protein (F) of paramyxo- and pneumoviruses, implying an evolutionary link between the two viral fusogens. Based on the structural information on the FV Env in two states, we propose a mechanistic model for its conformational change, highlighting how the interplay of its structural elements could drive the structural rearrangement. The structural knowledge on the FV Env now provides a framework for functional investigations such as the FV cell tropism and molecular features controlling the Env fusogenicity, which can benefit the design of FV Env variants with improved features for use as gene therapy vectors.

## Introduction

Foamy viruses (FVs) belong to the *Spumaretrovirinae* subfamily of the *Retroviridae* family [1]. They are unconventional retroviruses because their replication cycle displays several unique features that relate them to *Hepadnaviridae* and distinguish them from the better characterized members of the *Orthoretrovirinae* subfamily, such as the human immunodeficiency virus (HIV) and the human T-lymphotropic viruses [2, 3]. FVs are ancient viruses, estimated to have existed ∼450 million years ago [4–6]. This long coevolution with hosts might explain their seemingly non-pathogenic nature, which has been exploited for the development of FVs as vectors for gene therapy [7, 8]. FVs are prone to cross-species transmission, and humans can be persistently infected with simian FVs [2, 9].

As in all enveloped viruses, entry of FVs into host cells requires the fusion of viral and cellular membranes, a reaction catalyzed by the viral envelope protein (Env). Retroviral Env proteins belong to the class I of viral fusion proteins, whose members are also present in influenza viruses, filoviruses, paramyxoviruses, pneumoviruses, arenaviruses and coronaviruses. Class I fusion proteins share similar principles of oligomerization, proteolytic priming and activation (reviewed in [10, 11]). The general paradigm is that an N-terminal signal peptide (SP) directs their co-translational translocation into the endoplasmic reticulum, where they fold and oligomerize into trimers that remain membrane-anchored via a C-terminal transmembrane (TM) segment. The membrane-inserted SP is cleaved by a host signalase [12]. The protomers are further proteolyzed by an additional host protease to generate two subunits, an N-terminal, peripheral subunit (N-SU), which recognizes a cellular receptor or attachment factor, and a C-terminal, viral membrane anchored subunit (C-SU), which is maintained in a metastable, energy loaded state within the primed trimer [11]. Binding to a specific cellular receptor and / or protonation in the endosomal compartment [10, 13] triggers a major conformational change that drives membrane fusion. This process begins with the N-SU dissociation, which is in some cases released from the trimer, allowing the C-SU to spring out to form an extended intermediate [14–16]. In this state, the typically hydrophobic N-terminal end of the C-SU, termed “fusion peptide” (FP), inserts into the target membrane, while the protein remains anchored to the viral membrane via its TM segment [17]. The C-SU quickly folds back and collapses into a low-energy, post-fusion trimeric state bringing target and viral membranes into apposition and coupling the released free energy from protein refolding to the required dehydration of the two outer leaflets of the membranes to drive their fusion [18].

The hallmark of all class I fusion proteins in post-fusion state is a central trimeric coiled-coil that begins at the C-terminal end of the FP and is formed by a long *⍺*-helix that makes parallel interactions with its counterpart from the other protomers along the 3-fold molecular axis. The amino acid sequence in this region displays a seven-residue repeat pattern of non-polar residues termed “heptad repeat A” (HRA, or HR1) [19, 20]. The C-terminal segment, near the TM anchor, which is often also *⍺*-helical and can be identified by HRs in the amino acid sequence (termed “heptad repeat B” (HRB, or HR2)), packs in an antiparallel fashion along the interhelical grooves of the trimeric coiled coil, placing FP and TM anchors at the same end of a trimeric rod-like structure. The resulting six-helix bundle (6HB) is present in many class I fusion proteins in their post-fusion state [19–21].

FVs enter cells via receptor-mediated endocytosis [22]. Heparan sulphate has been identified as an important attachment factor for FV entry [23, 24]. The existence of a protein receptor has been postulated [25], but its identity remains elusive. Unlike the Env of most members of the *Orthoretrovirinae* subfamily, which are triggered by receptor binding at the plasma membrane, the FV Env mediated fusion occurs in the acidic environment of the endosomes [22]. Another distinguishing feature is the presence of two cleavage sites for proteolytic maturation in FV Env, an N-terminal furin-like site (residues 122-RRIAR-126) and a canonical furin site at position 567-RRKR-570 (Fig. S1A). Cleavage at the first site, mediated by cellular furin [26], separates the signal peptide from N-SU, while the second cleavage separates the N-SU from the C-SU. The SP remains associated with the mature FV Env trimer [26], which is composed of 3 subunits (SP, N-SU and C-SU), and has two membrane spanning segments, TM_1_ and TM_2_, within SP and C-SU, respectively (Fig. S1A). Previous studies of intact FV Env on viral particles provided low-resolution cryo-EM maps that informed about the general architecture of the molecule and revealed that Env trimers form a lattice on the virion surface [27]. The resolution of this structure was however insufficient to elucidate the atomic details of the protein or to understand its function.

Here, we describe the structures of the FV Env ectodomain determined by cryo-EM first in its post-fusion trimer of hairpins conformation, and then in the pre-fusion state. Our work shows that FV Env bears structural similarity to the fusion (F) protein of otherwise unrelated paramyxoviruses and pneumoviruses, with the notable insertion of an RBD upstream the furin site that separates N-SU and C-SU. Comparison of FV Env in its two conformations, pre- and post-fusion, allows a description of the functional transition of the protein to drive membrane merger, revealing insights into the mechanism used by this ancient fusogen.

## Results and discussion

### The post-fusion state

We expressed the predicted ectodomain of the gorilla FV Env protein (described in detail in Methods and Fig. S1) and determined its structure by cryo-EM at 3.1 Å resolution (Fig. S2, Table S1). The structure revealed a rocket shaped Env trimer in the post-fusion conformation characteristic for class I proteins, with a 6HB projecting from its base (Fig. 1). The RBDs are symmetrically arranged on the sides of a central shaft, with their long axes roughly parallel to the trimer axis (Fig. 1A). The central shaft bears striking resemblance to the fusion (F) proteins of paramyxo- and pneumoviruses (Fig. 1B) (see below), displaying the same head, neck, and stalk regions, formed by domains I, II and III as previously described for F [28, 29]. Domain I is a *β*-sheet domain, domain II has an immunoglobulin superfamily fold [30] and domain III has an *⍺*/*β* topology, featuring a prominent ‘core *β*-sheet’ (Fig. S3). When outlined on the primary structure of the protein, the domains of Env show a nested arrangement (Fig. 1C). The RBD, which constitutes a large portion of the N-SU, is the only domain made of a continuous stretch of amino acids (residues 217-543). The RBD is flanked by two linkers (L and L’), which connect it to domain III (Fig. 1D). Domain III is inserted between two *β*-strands that belong to domain I, which is in turn flanked by domain II elements.

**Figure 1:**
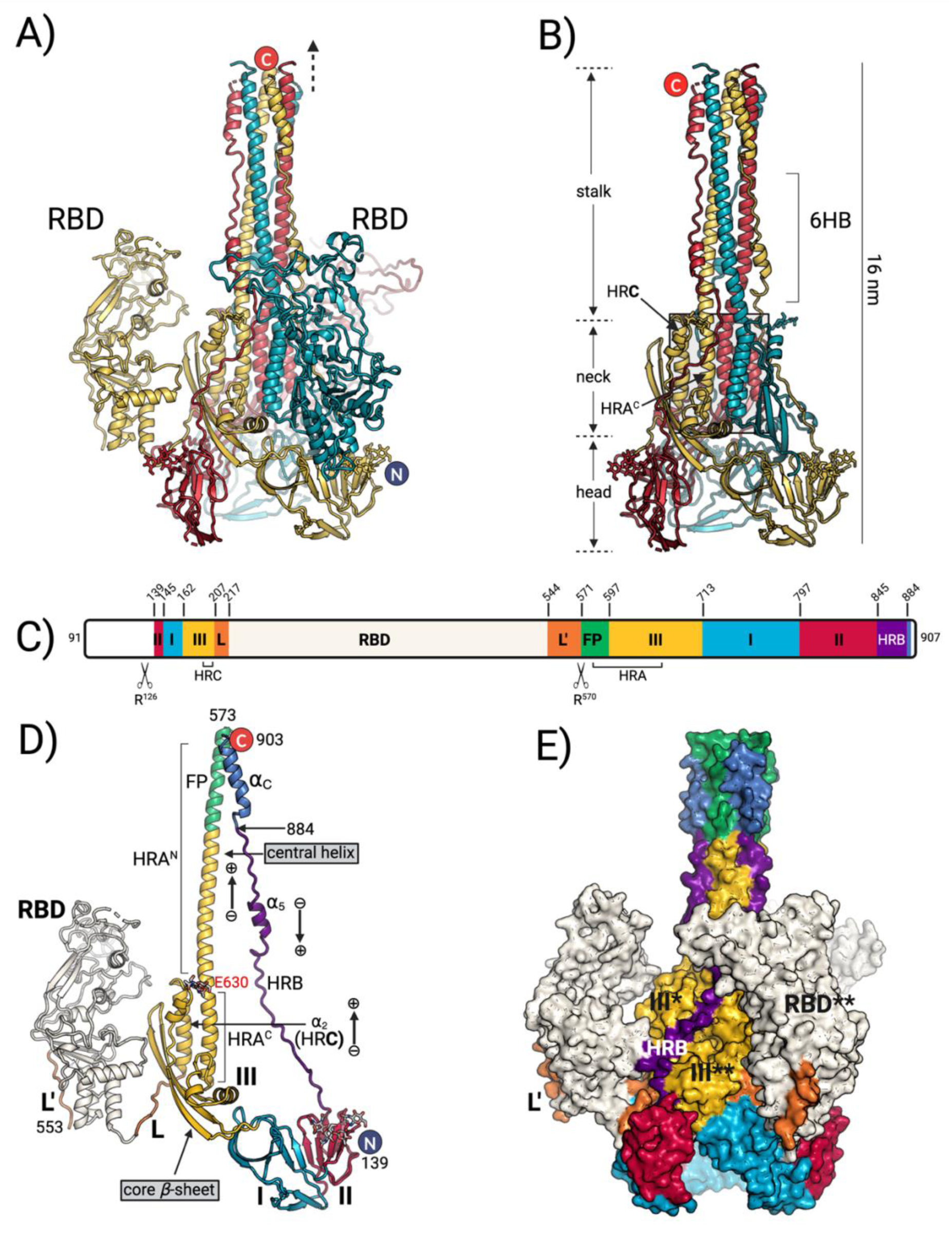
Post-fusion Env structure. **A)** Trimeric Env ectodomain is shown in cartoon representation, with each chain displayed in a different color. The first and last resolved residues (Glu 139 and Leu 903) are marked with letters N and C, respectively. Laterally positioned RBDs and the putative location of the membrane are indicated with RBD and dashed arrow, respectively. **B)** The central shaft (the trimeric Env ectodomain without RBDs) is shown in cartoon representation. The three HRA^C^ and three HR**C** *⍺*-helices, packing in parallel fashion, are indicated with the shaded box. **C)** Schematic representation of the Env ectodomain organization. The furin sites are indicated with the scissor icon. The domains are labelled with roman letters and colored as domain I (cherry red), domain II (azure) and III (bright gold). The first residue of each domain is indicated on the top. The other structural elements displayed are the fusion peptide (FP) (green), RBD linkers L and L’ (orange), RBD (beige), HRB (indigo) and C-terminal *⍺*_C_ helix (navy blue). The HR**C** (residues 186-204) and the HRA segments (residues 576-651) of domain III and indicated below the scheme. The terminal regions not resolved in the structure are shown in white (residues 91-138 and 905-907). **D)** The Env protomer is shown in cartoon representation with each domain colored according to the scheme in panel C). The central helix, composed of HRA^N^ and HRA^C^ segments, and the HR**C** helix are labelled. The arrows indicating the dipolar moment of *⍺*-helices (− and + corresponding to the C- and N-termini, respectively) are used to show the organization of helices. The core *β*-sheet (domain III) is marked. **E)** The trimeric Env ectodomain is colored by domain, according to panel C), and shown in molecular surface representation to illustrate the inter-protomer interactions. The star symbols (* and **) designate the neighboring protomers.

The FV Env coiled coil at the center of the 6HB is formed by a long *⍺*-helix (‘central helix’) that packs in a parallel fashion against its counterparts from the other protomers along the molecular 3-fold axis. The first resolved amino acid at the N-terminus of the central helix, Asn 573, lies only two residues after the furin site (residues 567-RRKR-570), which has undergone cleavage in our construct (Fig. S1B). The 21 turn-long central helix encompasses the HRA segment, and its N-terminal portion includes the region predicted to be the fusion peptide (FP) (Fig. S1C-D) [31, 32]. The 16 N-terminal turns of the central helix (termed HRA^N^ helix) coil around each other along the trimer axis, while its 5 C-terminal turns (termed HRA^C^ helix) run parallel to HRA^C^ of the other protomers and also pack against helix *⍺*_2_ of domain III (Fig. 1D). This helix *⍺*_2_, which was designated as “HR**C**” in F homologs (Fig. S3, Fig. 1B), connects to the RBD through a segment of random coil (linker L; residues 207-216) (Fig. 1D). The second linker, after the RBD (L’, residues 544-570), contains the furin cleavage site and is disordered in the structure.

At the C-terminal end of the construct, after domain II, the polypeptide chain adopts a random coil conformation, interrupted by *⍺*_5_ helix, and follows the trimer interface along the central shaft, interacting with domain III of the adjacent protomers (Fig. 1D-E). This segment is equivalent to the HRB linker and the HRB *⍺*-helix in the F proteins [33] (for simplicity we will apply the term HRB to designate the whole Env region spanning residues 845-884). The *⍺*_5_ helix packs in the grooves of the HRA^N^ coiled coil, giving rise to the 6HB. The last resolved residue, Leu 903, belongs to a C-terminal helix termed *⍺*_C_ (Fig. 1D) that is juxtaposed to the FP, the element presumed to interact with the target membrane during fusion (Fig. 1A).

The presence of the FP in the post-fusion form has not been seen in other class I fusion proteins, eother because it was disordered or because it was not included in the expression construct. The FP packs against the *⍺*_C_ helix (residues 886-902) at the tip of the coiled coil. The current paradigm is that an extended intermediate is formed during the fusion reaction, in which the FP interacts with the target membrane, while *⍺*_C_ remains in the proximity of the viral membrane. In the recombinant Env ectodomain, these two segments adopted a stabilizing conformation (Fig. S1E), which we think is an artifact since the interaction could not occur in the context of the full-length Env embedded in the lipid bilayer. Analysis of hydrophobic clusters at the interface between the coiled coil and the zippering HRB supports the idea, indicating that a stable 6HB would likely form even in the absence of the FP and *⍺*_C_ as the hydrophobic cluster formed by helix *⍺*_5_ is by far larger than the contacts keeping the *⍺*_C_ and FP together (Fig. S1D).

### The pre-fusion conformation

To stabilize the metastable pre-fusion state, the expression construct used to obtain the pre-fusion FV Env structure was modified to contain a C-terminal trimerization motif (foldon), mutations in both furin cleavage sites and an additional E630P mutation (described in detail in the Methods and Fig. S3). We calculated a cryo-EM map at a resolution of 3.8 Å that allowed tracing the polypeptide chain of FV Env (Fig. S5, Table S1). The obtained model was fitted in the 8.8 Å cryo-EM map reported for the full-length Env expressed on viral vector particles [27] with a correlation coefficient of 0.91, validating that the Env ectodomain structure represents the fold adopted by the protein displayed on virions (Fig. S4D).

The pre-fusion Env trimer is a barrel-shaped molecule with two large cavities along the central 3-fold axis, one at the top (membrane-distal) and one at the bottom (membrane proximal) side (Fig. 2A). The roof of the membrane-distal cavity is formed by the RBDs, which have a similar conformation to the one determined by X-ray crystallography [34]. The local resolution of the map in the region corresponding to the RBD loops is >5.5 Å, highlighting their flexibility even in the context of trimeric Env and implying a loose RBD-RBD interface (Fig. S5). The membrane-proximal cavity is closed off by domains I and II, and the *⍺*_5_ helix that packs underneath.

**Figure 2:**
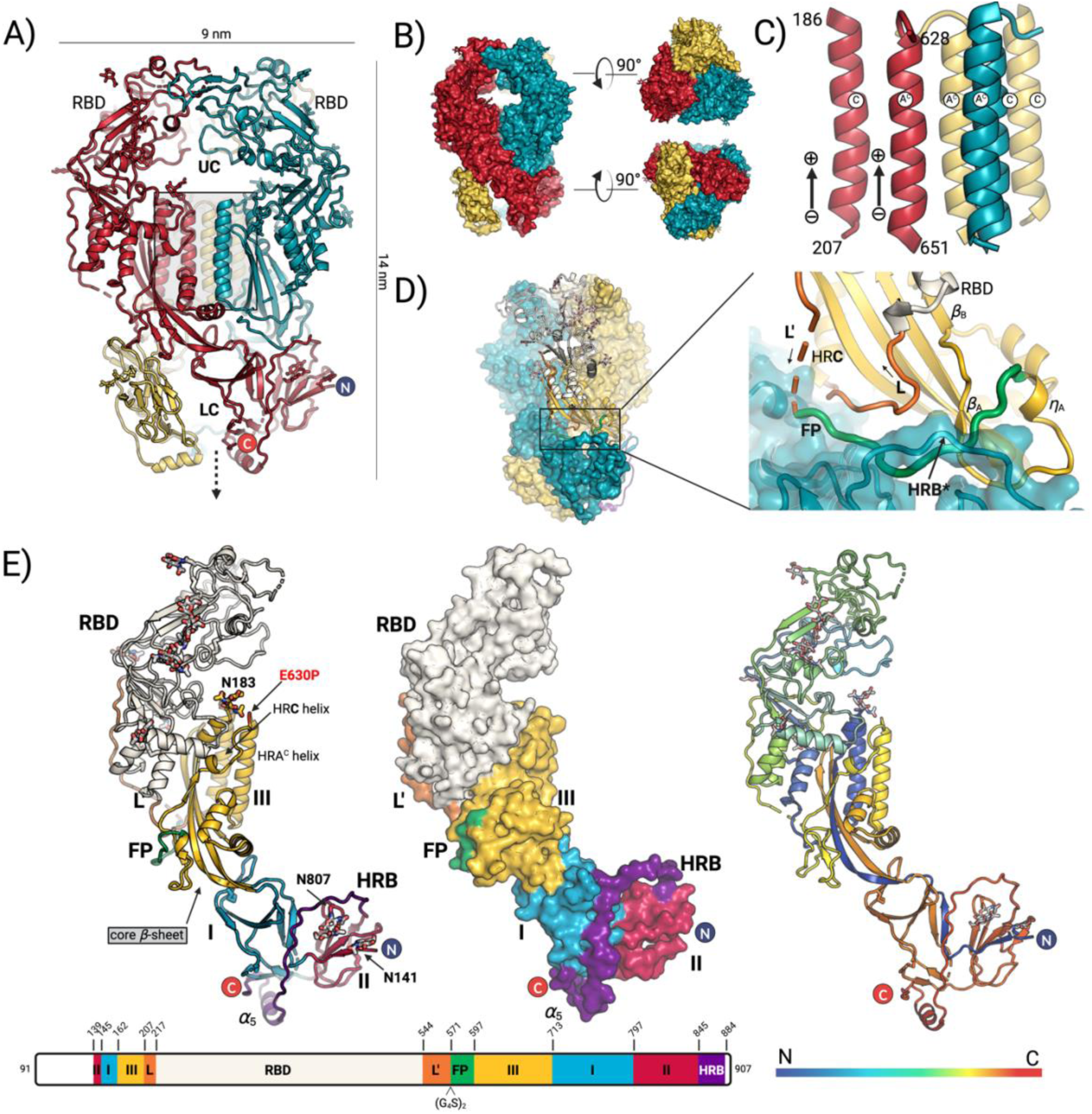
Pre-fusion Env structure. **A)** Trimeric Env ectodomain is shown in cartoon representation, with each chain displayed in a different color. The first and last resolved residues (Glu 139 and Ser 884) are marked with letters N and C, respectively. The putative location of the membrane is indicated with the dashed arrow. UC and LC designate the upper and lower cavity, respectively. **B)** Molecular surface representation of the trimer from panel A). **C)** Magnified region from the shaded box in panel A) is shown, representing the helical core comprised of 6 parallel *⍺*-helices (3 HRA^C^ (marked as ‘A^C^’) and 3 HR**C** helices (marked with ‘C’)). The arrows indicate the dipolar moments of *⍺*-helices (− and + corresponding to the C- and N-termini, respectively). **D)** Magnified region (right panel) of the area indicated with the black line box on the trimeric Env (left panel). One protomer is shown in cartoon and colored by domain. The two adjacent protomers are shown as molecular surfaces and are colored by chain (blue-green and yellow). The FP, which inserts between the linker L and the HRB* is labeled. The black arrows indicate the direction of the polypeptide chain. The structural elements in the HRA^N^ that follow the FP, the 3_10_ helix (*η*_A_) and strands *β*_A_ and *β*_B_, are marked. **E)** The Env protomer is shown in cartoon representation (left panel) with each domain colored according to the scheme in the panel below. The E630P mutation introduced to stabilize Env in its prefusion conformation (see Methods), is located between the HRA^N^ and HRA^C^ region and is marked with red letters. The molecular surface of the same protomer is shown in middle panel, illustrating the wrapping of the HRB around domains I and II, ending the with the *⍺*_5_ helix at the bottom of the molecule. The cartoon representation of the Env protomer is colored in blue to red spectrum, corresponding to the N- to C-termini in the right panel.

The three Env protomers wind about each other (Fig. 2B), making tight contacts in the central region through the HRA^C^ and the HR**C** (*⍺*_2_) helices that run parallel to each other (Fig. 2C). The resulting helical core separates the membrane-distal and proximal cavities (Fig. 1B). The segment corresponding to the HRA^N^ helix in the post-fusion form, which includes the FP, adopts essentially a random coil conformation, with short segments of secondary structure (a 3_10_ helix *η*_A_, followed by strands *β*_A_ and *β*_B_ and a short *⍺*_A_ helix (Fig. S3, Fig. S6)). The FP runs below linker L of the same protomer, settling into a crevice formed by the HRB* segment, which partially secludes the FP from the outside (Fig. 2D) (the star sign * is used throughout the text to indicate belonging to the neighboring protomer). The stabilizing E630P mutation is in a loop connecting the HRA^N^ and HRA^C^ helices (Fig. S6, Fig. 2E). Residues 557-581, comprising the second half of L’ are disordered, indicating that the furin site, which was substituted by a linker in our construct, would be exposed on the sides of the Env trimer where it is accessible for cleavage. The HRB region adopts an extended polypeptide conformation that wraps around domains II and I, ending with a short *⍺*_5_ helix and strand *β*_17_, which augments the domain I *β*-sheet (Fig. S3). The last amino acid with clear density, Ser 884, is immediately upstream the C-terminal helix *⍺*_C_, which was observed only in the post-fusion form even though the expression constructs for the pre- and post-fusion Env ectodomains had the same boundaries (Fig. 1).

Several glycosylation sites were resolved in the FV ectodomain. The glycan attached to Asn 183, in a loop connecting helices *⍺*_1_ and *⍺*_2_ (HR**C**) of domain III (Fig. 2E), partially fills the membrane-distal cavity. The six glycans present in the RBD at Asn 311, 346, 373, 390, 404, and 524, project from the exposed surface of the trimer and do not penetrate into the cavity. Two additional glycans are visible in the structure, located on domain II at the base of the trimer, and attached to Asn 141 and Asn 807 (Fig. 2E).

Out of 20 cysteine residues, 18 were found to form 9 disulfide bonds (DSB). C562 could not be modelled due to poor quality of the cryo-EM map in this region, which includes the N-SU linker L’, leaving C751 from the C-SU domain I unpaired. An unsharpened cryo-EM map provided the hint that C562 and C571* were in position to make an inter-protomer disulfide bond (Fig. S7A). SDS-PAGE of cleaved and uncleaved constructs under non-reducing conditions showed that the Env trimer is covalently linked by disulfide bonds between the N-SU from one protomer and the C-SU* (see Methods, (Fig. S7B, C)). The inter-protomer DSB could be only attributed to C562-C751*, since all the other cysteine residues are engaged in intra-protomer disulfide bonds. Analysis of a C562A mutant confirmed this prediction (Fig. S7C).

### Additional insight into pre-fusion FV Env from AlphaFold Multimer predictions

To obtain structural insights on regions that were not included in the expression construct or were not resolved in our stabilized Env ectodomain, we performed *ab initio* structural prediction of full-length FV Env using Alphafold multimer (AFM) [35]. The AFM model for the full-length Env trimer has the predicted template modelling (pTM) score of 0.77 (the pTM reflects the expected similarity between the predicted and the real structure), and the interface pTM (ipTM) score of 0.76 (indicates the confidence of the docked protein interface domains). The global LDDT value for the AFM model is 66.75, and the ‘LDDT per residue’ values are >70 for majority of the ectodomain residues, except for the exposed loops containing the two cleavage sites and the FP segment (Fig. 3A, Fig. S8A). Although the relative orientation of the domains I and II within each protomer varies in comparison to the experimental structure (Fig. 3A), the predicted trimeric model is strikingly similar to the experimentally determined structure, as indicated by the room mean square deviation (rmsd) of 1.7 Å for the superposition of 378 C_*⍺*_ atoms of the domain III (residues 162-206 and 631-712). The folding of the individual domains is preserved, as indicated by rmsd values <1.4 Å (Table S2).

**Figure 3:**
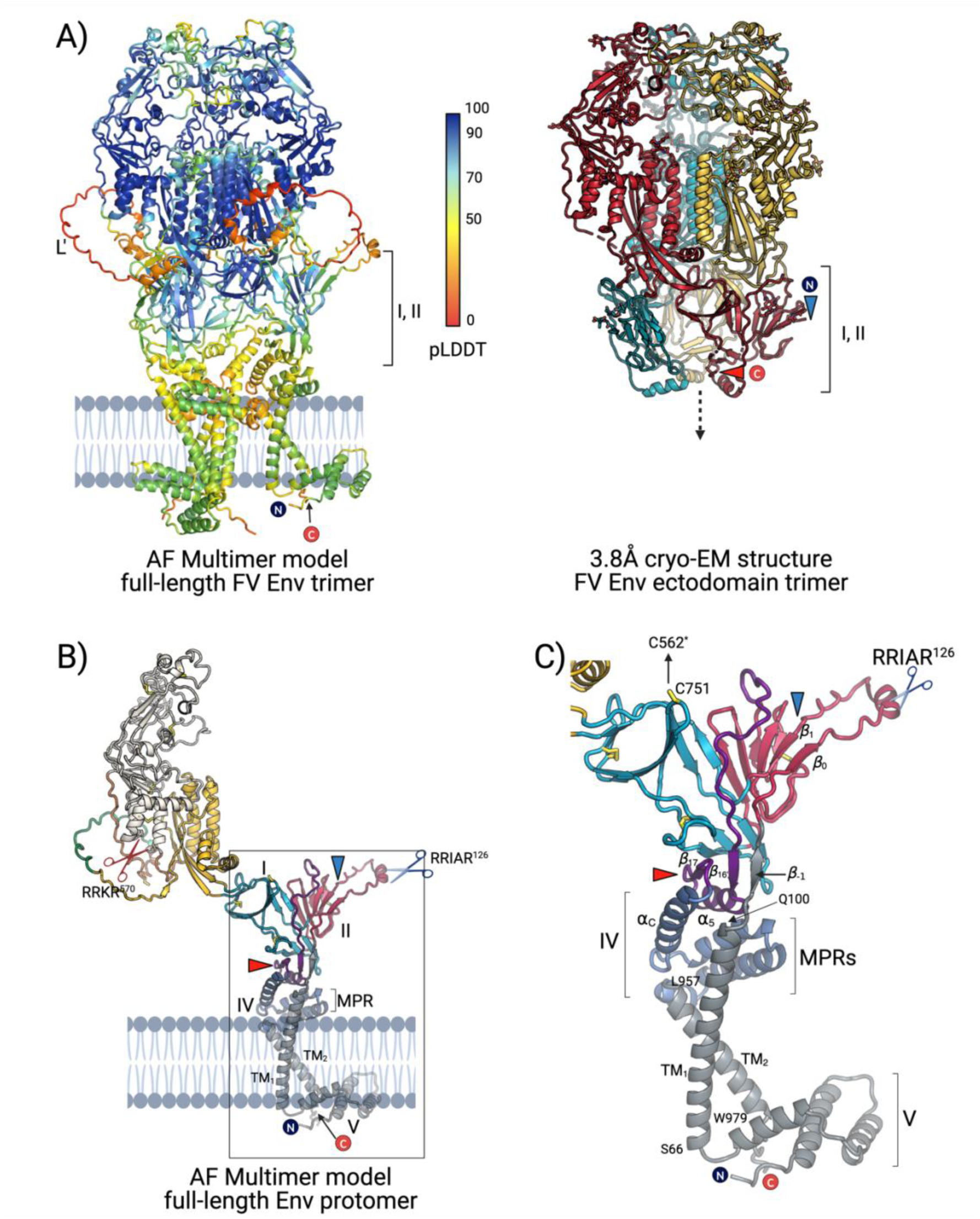
AF Multimer predicted model of the full-length Env trimer in the pre-fusion state. **A)** The AFM model of the full-length trimer is colored according to the per residue confidence metric called ‘predicted local distance difference test’ (pLDDT). The pLDDT can have a value between 0 and 100, with the higher pLDDT number the higher the confidence for the residue placement in the model. The predicted location of domains I and II, upwards compared to the experimental structure, is indicated, as well as the N- and C-termini. The experimental cryo-EM structure of the pre-fusion Env ectodomain is shown for comparison on the right. The protomers are each shown in a different color. The blue and red triangles designate the first and last resolved residue, Glu 139 and Ser 884, respectively. **B)** A protomer of the AFM model shown in A) is displayed and colored by domain using the color scheme defined in Figure 1C. The domains are marked with roman letters I-V. The helical bundle that comprises *⍺*_C_ and the putative MPR helices is colored in navy blue. The TM helices belonging to the SP and C-SU, the TM_1_ and TM_2_ respectively, are indicated and colored in grey together with the helical bundle of the luminal side that forms domain V. The blue and red triangles designate the first and last resolved residue of the experimental structure shown in panel A). **C)** The membrane proximal region of the Env pre-fusion model shown in panel B) is magnified to indicate the packing of domain IV *⍺*-helices constituting putative MPRs, and *β* strands preceding the furin-like cleavage site. The side chain of C571, which forms an inter-protomer DSB with C562* in the AFM model is shown in stick representation.

AFM prediction revealed that the SP residues 103-106 and 112-116 form two *β*-strands, termed *β*_-1_ and *β*_0_, respectively. Strand *β*_-1_ runs antiparallel to another predicted strand (*β*_16_’, residues 862-865), which belongs to the HRB region and adopts a random coil conformation in our experimental structure (Fig. 3B). It is possible that the interactions between these two *β*-strands further stabilize the HRB in the pre-fusion conformation by tying together the Env segments close to N- and C-termini. The *β*_0_ strand (residues 112-116) forms a *β*-hairpin with strand *β*_1_, which was resolved in our structure, projecting the SP furin-like site to the side of the trimer and augmenting a domain II *β*-sheet (Fig. 3, Fig. S3). When placed in the reported cryo-EM map of full-length Env [27], the *β*_0_-*β*_1_ hairpin fills the density that connects different trimers on the surface of viral particles, indicating that it could be an important structural element for the organization of the observed Env lattice (Fig. S8B).

AFM further predicts that the C-terminal helix *⍺*_C_, which was not observed in the experimental structure of pre-fusion Env, is part of an *⍺*-helical bundle, termed here domain IV, at the bottom of the pre-fusion trimer (Fig. 3B). This bundle contains 4 *⍺*-helices, and closes the lower cavity of the Env trimer, reinforced in part by the long TM_1_, which crosses with the C-terminal TM_2_ anchor at an angle (Fig. 3B). The side chains of strictly conserved His 906, predicted to be in a loop downstream of *⍺*_C_ helix, pack against each other at the 3-fold axis (Fig. S8B), suggesting that protonation of His 906 could destabilize the pre-fusion trimer in the acidic endosomal environment. Furthermore, two amphipathic *⍺*-helices of the domain IV bundle (*⍺*_6_ and *⍺*_7_; Fig. S6) are predicted to pack together roughly at the same height as the N-terminus of TM_2_ (residue Q100), in an orientation parallel to the viral membrane and facing the membrane with their hydrophobic sides (Fig. 3C). This observation suggests that the domain IV *⍺*-helices may play the role of membrane-proximal regions (MPR), which can be involved in membrane destabilization during fusion (reviewed in [36]). Poor density was observed at the bottom of the maps of the pre-fusion Env ectodomain construct (*ab initi*o models, classes 0 and 3, Fig. S5) suggesting mobility of domains I and II, possibly due to the absence of the MPRs which were not included in our expression construct to avoid solubility problems. Finally, AFM also predicts another *⍺*-helical bundle in the cytosolic side of the viral membrane (domain V) (Fig. 3C, Fig. S3).

### FV Env and paramyxo- and pneumovirus F share a common structural core indicating their evolutionary relatedness

The topological distribution of secondary structure elements in FV Env domains I, II and III (Fig. S3C) shows only minor variations to those observed in paramyxovirus and pneumovirus F [37, 38]. The rmsd calculated for superposition of 147 C_*⍺*_, corresponding to domains I, II and III of FV Env and F of parainfluenza virus type 3 is 3.6 Å, indicating comparable three-dimensional organization of the 3 domains (Fig. 4). The intertwined pattern of nested domains shown in Fig. 1C was first described for paramyxovirus and pneumovirus F (Fig. S9) [38, 39], and was also observed, along the structural similarity, for the fusion subunit of the coronavirus Spike protein extending the homology (Spike does not have domain II and domain I is smaller) [40, 41]. It is the same structural elements, the core *β*-sheet and the helical bundle that bear structural similarity in F and Spike, and in Env. The conservation of the core in FV Env now provides an additional evolutionary link between paramyxo-, pneumo- and coronaviruses, with the *Spumavirinae* subfamily of the retroviruses.

**Figure 4:**
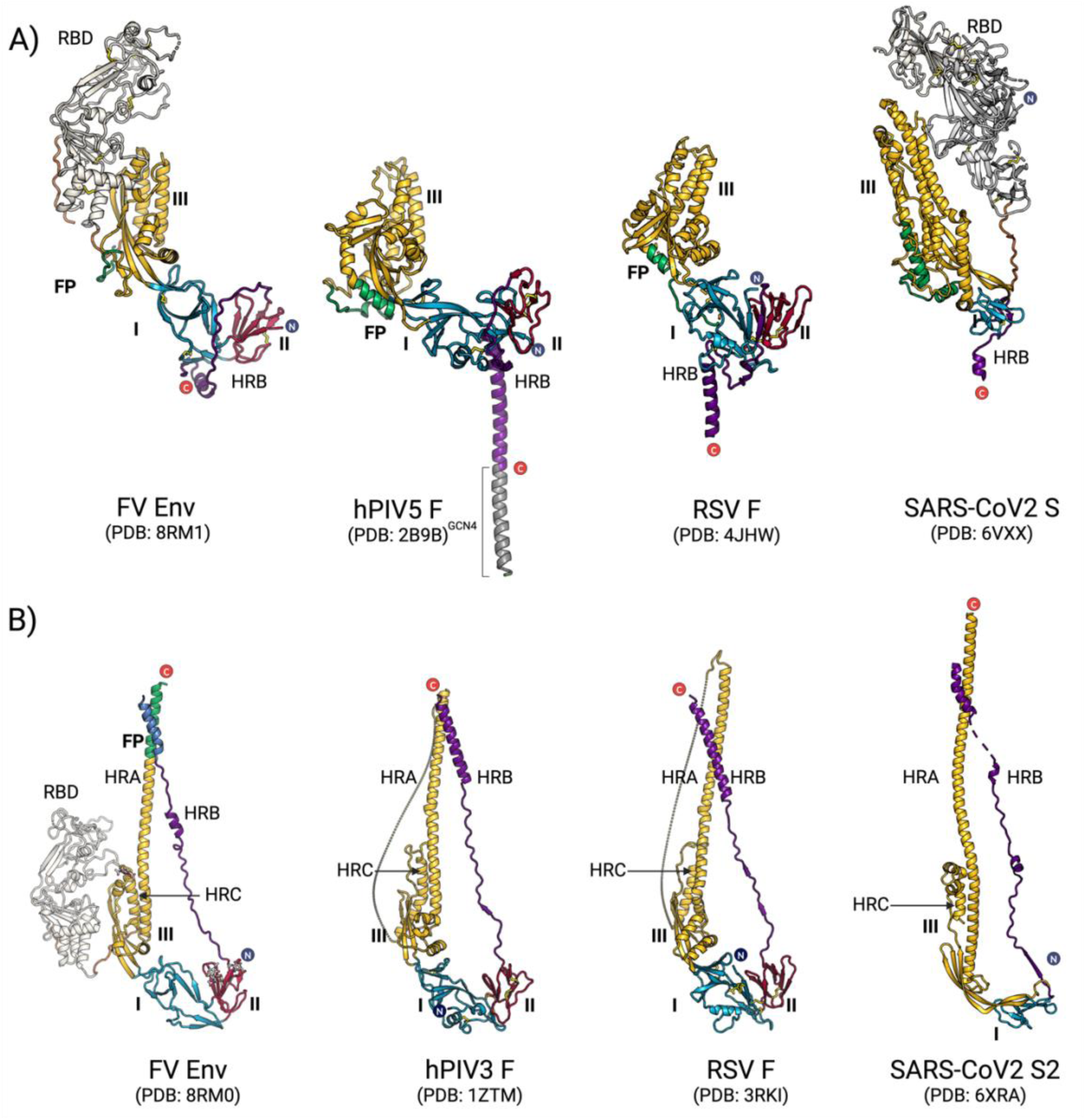
Structural similarity between fusion proteins of FV, paramyxo-, pneumo- and coronaviruses. Pre-fusion and post-fusion structures are shown on panels **A)** and **B)** respectively. Only protomers are displayed for clarity. Domains are colored according to the scheme in Figure 1C. hPIV stands for human parainfluenza virus (now called human respirovirus, member of *Paramyxoviridae*), RSV for respiratory syncytial virus (now called human orthopneumovirus, member of *Pneumoviridae*), and SARS-CoV 2 for severe acute respiratory syndrome coronavirus 2 (member of *Coronaviridae*). PDB codes for each structure are indicated. The terminology used for coronavirus Spike domains [70] is different than one established for F proteins; for simplicity we applied the former to all the proteins belonging to this subset.

The sugar attached to highly conserved Asn 183 of FV Env (Fig. S6), which projects into the membrane-distal cavity and partially covers the HRA^N^ segment in the pre-fusion conformation, does not form polar contacts with the protein and does not have an extensive buried surface area (Fig. 2E), suggesting a role other than a structural one, which we attributed to the oligosaccharide linked to the RBD Asn 390 [34]. The F protein of some of pneumo - and paramyxoviruses contains a glycan at the equivalent location, i.e. just upstream of HR**C** helix, which is fully exposed at the apex and was shown to be important for immune evasion [42, 43]. We speculate that the glycan at Asn 183 could have a similar role in the FV Env, masking potential neutralizing epitopes that are not occluded by the RBDs, or the epitopes that may become exposed during RBD movements.

Similar to the structure of the pre-fusion uncleaved paramyxovirus F protein [39], the FP of FV Env wraps for the most part around the exterior of the trimer at the inter-protomer interface [39]. In contrast, in the pre-fusion structure of pneumovirus F, the FP was found packing against an adjacent protomer but buried within the interior of the F trimer [38]. A unique feature that distinguishes pneumovirus F from its paramyxovirus and FV counterparts is the presence two consecutive furin sites preceding the FP (Fig. 4), both of which must be cleaved for the pre-fusion trimer to assemble, as the intervening p27 segment interferes with trimerization [44].

The most prominent feature that distinguishes Env from protein F in the pre-fusion form is the RBD insertion in Env, in line with FVs not having a dedicated, additional protein for receptor binding, as paramyxo- and pneumoviruses do [45]. In this context, the N-SU and C-SU of FV Env, termed also the surface (SU) and TM subunits, are equivalent to the F subunits F2 and F1, respectively, separated by the amino acidic sequence of the RBD that is accommodated, in the tertiary structure, at the periphery of the F scaffold (Figure 1).

Another important difference between the pre-fusion conformations of F and FV Env is observed at the C-terminal region. The HRB segment connecting domain II to the *⍺*_5_ helix of Env initially follows a path similar to that of its counterpart in the F proteins, interacting along the sides of domain I to reach the “bottom” of the trimer. However, the HRB helix *⍺_5_* packs underneath domain II in an orientation parallel to the viral membrane in Env (Fig. 2E, Fig. 5B), rather than forming the coiled coil ‘stalk’ structure observed in paramyxovirus F [39] and coronavirus Spike [46], or the inverted pyramid 3-helix bundle observed for pneumovirus F [38]. This difference may arise from constrains imposed by two membrane spanning segments in the FV Env. TM_1_ is required because the preceding N-terminal region, interacts with the Gag protein and is essential for FV budding [47, 48], in contrasts to orthoretrovirus Env, which binds to Gag via the cytosolic C-terminal segment downstream of TM_2_.

**Figure 5:**
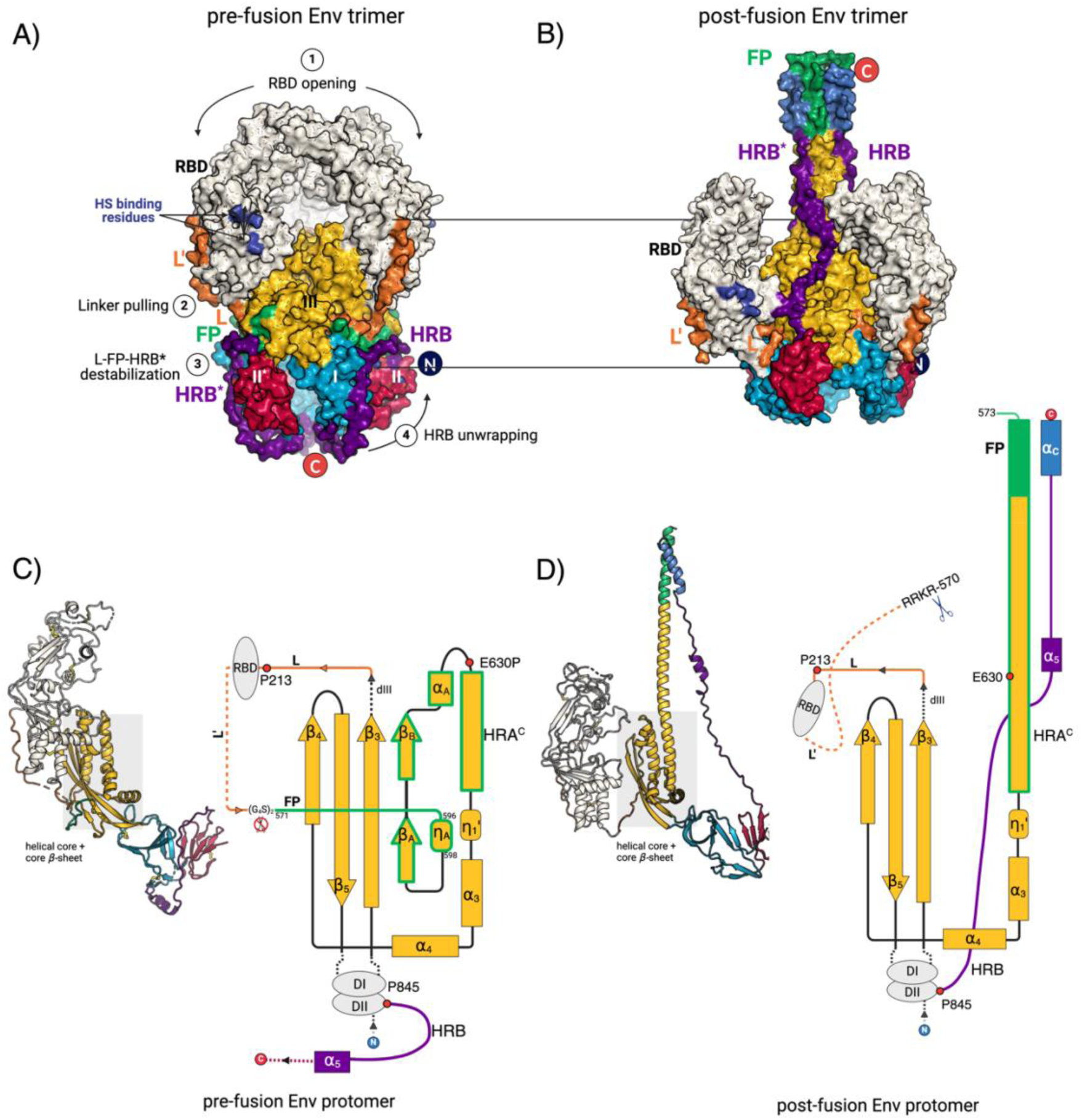
Insights into the conformational change mechanisms from the comparisons of pre- and post-fusion Env structures. **A)** and **B)** are molecular surface representations of the pre-fusion and post-fusion Env trimer, respectively, colored by domain and according to the scheme from Figure 1C. The pre- and post-fusion structure were aligned on the domain III helical core (HRA^C^ and HR**C** helices). The heparin sulphate (HS) binding residues in the RBD domain, K342, R343, R356 and R369, are highlighted in dark blue. Numbers 1-4 illustrate a possible order of events during the conformational change: 1-lateral movements of the RBDs, 2 – RBD pulling of the linker L, 3 -destabilization of the L-FP-HRB* interface and liberation the FP, and 4 – unwrapping of the HRB, which flips and zippers down the post-fusion coiled coil. Star symbol * designates the structural elements belonging to the adjacent protomer. **C)** and **D)** are the cartoon representations of the Env protomers in the pre- and post-fusion state, respectively. The inlets show the secondary structure topology of the domain III elements *β*-core sheet and the HRA region that undergoes refolding, and which is highlighted with green borders. The locations of the strictly conserved prolines, residues 213 and 845, which are the points where the polypeptide chain changes direction, are indicated with red circles.

### The proposed model for the FV Env conformational change

During the conformational transition, the RBDs undergo a major repositioning (their centers of mass shift ∼30 Å, Table S3), while maintaining their fold (Table S4). As was observed for the F proteins, the FV Env domains I and II also preserve their 3D folds and relocate as rigid bodies (Table S4; Fig. 5). The core *β*-sheet and HR**C** helices from domain III remain unchanged in the Env, F and Spike, and serve as a hub around which the HRA segment refolds. The helical core however collapses inwards during the pre- to post-fusion transition in the F proteins, compacting the head region [38, 39]. Conversely, the conformation of the helical core is maintained in FV Env (superposition of C_α_ atoms has the rmsd of 1.2 Å), conferring a larger core around which the structural rearrangement takes place.

In the pre-fusion FV Env state, the RBDs are held in place by loose inter-protomer interactions at the top of the trimer and by contacts between linker L, which precedes the RBD, with the FP and HRB* segment (Fig. 5A, Fig. 2D). Based on the comparison of the pre- and post-fusion structures of FV Env, we propose the following sequence of events for the conformational transition. Binding to HS and/or a putative receptor pulls the RBDs away from each other, resulting at least in a partial opening of the Env trimer. The movement of the RBDs, and consequently the linker L it is attached to, destabilizes the interface between L, FP and HRB* (Fig. 1D), liberating the two latter segments to establish new contacts. As a result, the HRB region unwraps from domain I and swings out to extend upwards, pulling along domains II and I. Complete dissociation of the RBD trimer is necessary to create space for the refolding of the HRA^N^ segment into the central helix to form the coiled coil and project the FP into the target membrane to make an elongated intermediate. The acidic environment of the endosomes and the residues sensitive to pH, such as the strictly conserved His 906 (Figure S8B), could play a role in destabilization of the MPRs and TM anchors, which also need to separate for membrane fusion to complete.

### Concluding remarks

Unlike class II and III fusogens, the fold of class I fusion proteins cannot be considered to imply homology, because the simple coiled coil motif, accommodating the C-terminal segments of the protein in the interhelical grooves, could have evolved independently more than once during evolution. We propose that the class I proteins can be divided into at least in two sub-classes of likely different origins. One subclass, which we term class Ia, includes orthoretrovirus Env, influenza virus HA, filovirus and arenavirus GP, where the membrane fusion function is strictly confined to the C-SU that results from the maturation cleavage. In the paramyxo- and pneumovirus F, coronavirus Spikle, as well as FV Env, the membrane fusion protein is not ascribed to the C-SU, but to the N-SU/C-SU heterodimer, implying a likely independent evolution scenario between the two subclasses. Despite the lack of amino acid sequence conservation, the subclass Ib fusion proteins from paramyxoviruses, pneumoviruses, coronaviruses, and FVs share a common structural core indicating they derive from a distant common ancestor (Fig. 4). Within subclass Ib, the F protein is the smallest, with a fold that represents the minimal structural scaffold that can accommodate receptor binding modules, either at its N-terminus, as in coronavirus Spike, or inserted between the two subunits, as in the FV Env (Fig. 6, Fig. S9). Phylogenetic analyses revealed that FVs originated > 450 million years ago making them one of the most ancient group of viruses known so far [6]. One could speculate that the FV *env* could have provided a template for excisions or insertion of fragments, such as the RBD. Consistent with this idea are the findings highlighting the modular nature of simian FV genomes and existence of two *env* variants that differ in a region that is flanked by two recombination hotspots within the N-SU [49].

**Figure 6:**
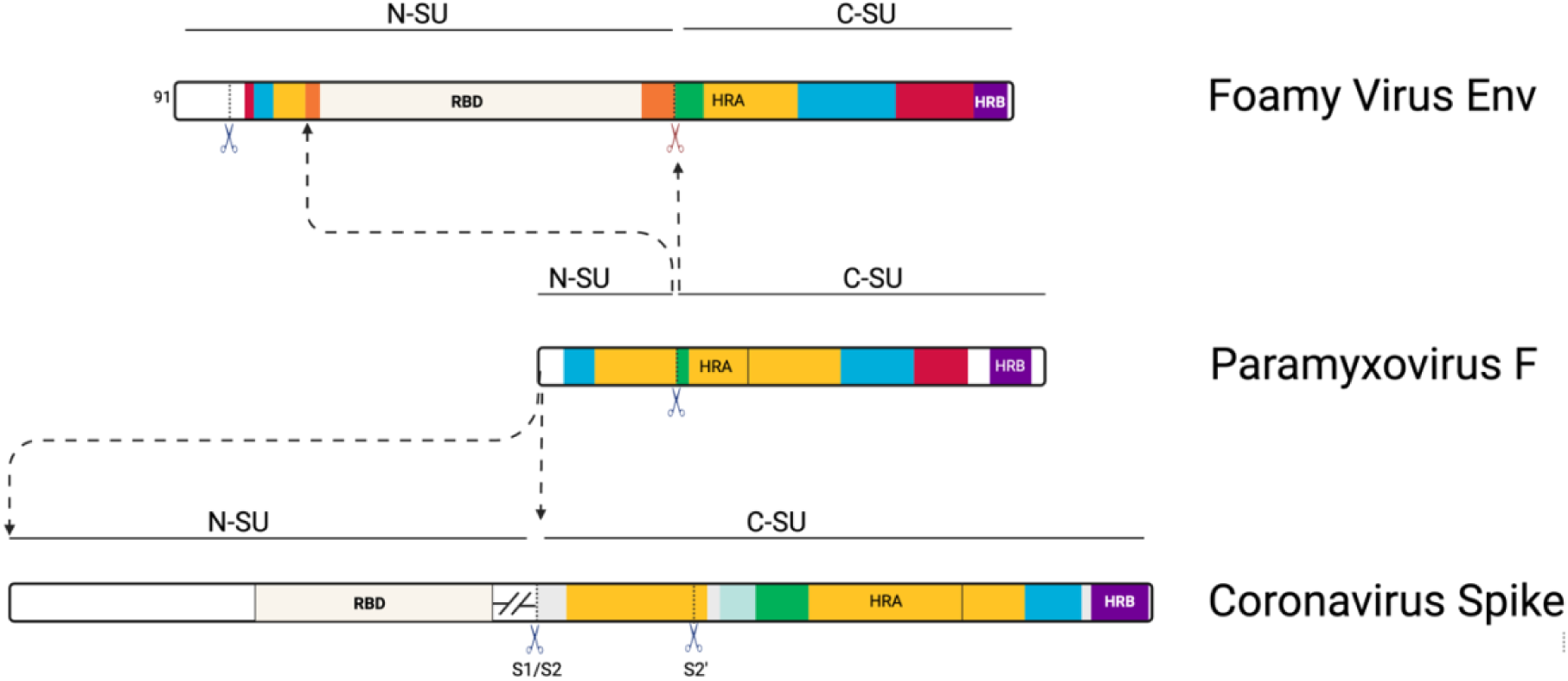
Modular nature of class Ib fusion proteins. Schematic representations of primary structure of the fusion proteins from FVs, paramyxoviruses and coronaviruses. The segments constituting domains I, II and III are colored according to the scheme from Fig. 1C. The dashed arrows indicate the incorporation sites of the RBD modules. The furin cleavage sites are marked with the scissors symbol. The light green box in coronavirus Spike, just upstream of the FP, corresponds to the FP proximal region, segment observed only in the Spike [71] (see Fig. S9 for details).

The lack of structural similarity between the FV Env and the Env of orthoretroviruses, along with the reported novel fold for the FV RBD [34] further support the classification of FVs as a separate retroviral subfamily, which had originally been done based on the unique features of their replication cycle [1]. Many questions however remain to be answered, primarily the requirement and nature of the cellular receptor and the fusion triggering mechanism that may rely on the presence of receptor as well as acidification. The atomic-resolution structures of the FV Env in the pre- and post-fusion states now provide a framework for further functional investigations such as the FV cell tropism and molecular features controlling the Env fusogenicity. The structural information can also be exploited for design of FV Env variants with improved features for their use as gene therapy vectors.

## Methods

### Construct design

We predicted the boundaries of the Gorilla Simian Foamy Virus Env ectodomain (GII-K74 strain, accession number JQ867464) using Phyre2 [50], which predicted two transmembrane helices: the first between residues 60 to 90, and the second one between 948 and 978. We tested constructs bearing C-terminal truncations at residues 907, 936 or 945 and selected 91-907 as the best ectodomain candidate based on its higher expression.

The WT Env ectodomain served as the template for introduction of mutations commonly used to stabilize viral fusion proteins in their prefusion conformation [51]: 1) R126A mutation in the furin-like site on the leader peptide prevented the SP/N-SU cleavage [26], 2) the canonical furin site (567-570) was replaced with a flexible and non-cleavable linker (GGGGSGGGGS) abrogating the cleavage, 3) foldon trimerization motif was added at the C-terminus (SAIGGYIPEAPRDGQAYVRKDGEWVLLSTFLG). The resulting expression construct was termed variant 1 (V.1 on Fig. S4A). Cryo-EM analysis and the 3D *ab initio* models revealed that about 40 % of the particles appeared to have a conformation reminiscent to the one obtained by docking the RBDs into a low-resolution reconstruction of the pre-fusion full-length Env trimer on virions [27, 34]. This conformation is characterized by having the RBDs - which are readily identified by their characteristic bean shape - assembled near to each-other, in a closed conformation (Fig. S4B). To further stabilize the pre-fusion form, we performed a proline-scan around a region of central helix where we anticipated a helix break in the pre-fusion form, based on similar work on other class I fusion proteins [52, 53]. We thus identified mutant E630P (construct V.2 on Fig. S4A), which resulted in ∼70% of the particles appearing in the closed, presumably pre-fusion form, and which allowed us to generate maps and build an atomic model. The E630P did not have an effect on vector particle production, infectivity and cell binding (Fig. S4D), indicating that the mutation itself was not sufficient to lock the Env in the pre-fusion state, and suggesting a larger importance of the furin site substitution and foldon for Env stabilization.

### Protein expression and purification

The codon optimized DNA sequence coding for the wild type gorilla FV Env ectodomain from strain SFVggo_huBAK74 (accession code JQ867464.1, [54]) was synthesized by Genscript and cloned into a modified pMT/BiP insect cell expression plasmid (Invitrogen) designated pT350, which contains a divalent-cation inducible metallothionein promoter, the BiP signal peptide at the N-terminus (MKLCILLAVVAFVGLSLG), and a double strep tag (DST) (AGWSHPQFEKGGGSGGGSGGGSWSHPQFEK) at the C-terminus [55]. Mutagenesis to obtain the stabilized ectodomain construct, as well as the R570T and C562A mutants, was performed by Genscript.

Each plasmid was co-transfected in Drosophila Schneider line 2 cells (S2) with the pCoPuro plasmid for puromycin selection [56]. Both cell lines underwent selection in serum-free insect cell medium (HyClone, GE Healthcare) containing 7 μg/ml puromycin and 1% penicillin/streptomycin. For the protein production stage, the cells were grown in spinner flasks until the density reached ∼1 × 10^7^ cells/ml, at which point the protein expression was induced with 4 μM CdCl_2_. After 6 days, the cells were separated by centrifugation, and the supernatants were concentrated and used for affinity purification using a StrepTactin column (IBA). The proteins were further purified by size exclusion chromatography (SEC) on a Superose6 16/60 (Cytiva) column (wild type ectodomain) or using a Superdex 200 10/300 (Cytiva) column (stabilized ectodomain) in 10 mM Tris, 100 mM NaCl, pH 8.0. Eluted fractions were analyzed by SDS-PAGE under reducing and non-reducing conditions, and some of them were pooled and concentrated (wild type ectodomain) or stored separately (stabilized ectodomain).

### Sample preparation for cryo-EM

To find the best construct stabilized in the prefusion conformation, a screening of candidates was performed by preparing grids with 3 μl of protein (1.5 µM trimer in SEC buffer), which were applied on Quantifoil R1.2/1.3 200 mesh copper grids that been glow-discharged twice using a Pelco glow discharge system at 15 mA for 25 s. Samples were immediately vitrified in 100% liquid ethane using a Mark IV Vitrobot (Thermo Fisher Scientific) by blotting for 3.5 s with Whatman No. 1 filter paper at 9°C and 100% relative humidity. 3 μl of the selected stabilized ectodomain were added to Quantifoil R2/2 200 mesh copper grids with a 2 nm continuos carbon film on top (Delta microscopies), which were glow-discharged twice beforehand using a Pelco glow discharge system at 15 mA for 25 s. Samples were immediately vitrified in 100% liquid ethane using a Mark IV Vitrobot (Thermo Fisher Scientific) by blotting for 3.5 s with Whatman No. 1 filter paper at 9°C and 100% relative humidity, after 10-s waiting time. 3 μl of the wild type ectodomain (1.0 µM trimer in SEC buffer) were added to Quantifoil R1.2/1.3 200 mesh copper grids (Delta microscopies), which had been glow-discharged as indicated before. Samples were immediately vitrified by blotting for 4 s with Whatman No. 1 filter paper at 9°C and 100% relative humidity.

### Cryo-EM data collection, processing, refinement and modelling (post-fusion form)

Data of the wild type ectodomain were acquired on a Titan Krios transmission electron microscope (Thermo Fisher Scientific) operating at 300 kV, using EPU automated image acquisition software (Thermo Fisher Scientific). Movies were collected on a Gatan K3 direct electron detector operating in counting mode at a nominal magnification of 105,000× (0.85 Å/pixel) using a defocus range of −1.0 to −3.0 µm. Movies were collected over a 2s exposure and a total dose of ∼50 e−/Å^2^.

Data were processed by using Relion-3.0 [57] following the workflow shown in FIG S2. In brief, row movies were processed using MotionCor2, with a five-by-five patch-based alignment, keeping all frame and dose-weighting up to the total exposure. The CTFs of the dose-weighted images were determined using CTFfind-4.1 [58]. Images having a CTF maximal resolution worse than 4 Å, as well as those with important ice contamination or large carbon area, were discarded. Particles were picked using the Relion Laplacian of Gaussian, they were extracted and submitted to three rounds of 2D classification. The particles from the selected classes were used to generate an initial 3D model, which in turn was employed to perform 3D classification. The position of particles from one class were refined, the resulting map (4 Å resolution) was filtered to 20 Å and used to perform a round of auto-picking with a 3D reference. The new set of particles was extracted, used for two cycles of 2D classification and the those from the best classes were submitted to a 3D classification job with 50 iterations and the initial 3D model as reference. Particles belonging to the best 3D class were re-extracted (without binning) and used to perform per-particle CTF refinement, particle polishing, and another per-particle CTF refinement (shiny particles). A final round of 3D classification rendered four classes, two of which were selected to combine their particles in a last 3D refinement round, resulting in a 3.1 Å resolution map.

Building of the N-SU atomic model started by placing the crystal structure of the RBD (PDB: 8AIC, [34]) into the sharpened EM map using Chimera and manually extending the N-terminus in Coot. The first residues of the TM atomic model were built by the map-to-model tool from the Phenix suite, which generated a segment of the HR1 helix that was manually extended in N- and C-terminal directions in Coot. Following this strategy an initial model was obtained and it was refined with one round of real space refinement including simulated annealing in Phenix, followed by iterative rounds of manual building and real-space and B-factor refinement in Coot [59] and Phenix, with secondary structure restraints. As before, symmetry operators were obtained from the EM map and they were used to place three copies of the protomer within the map using tools from Phenix. Validation of model coordinates was performed using MolProbity [60].

### Cryo-EM data collection, processing, refinement, and modelling (pre-fusion form)

Screening of the stabilized ectodomain candidates was performed by collecting Single-particle cryo-EM data on a Glacios transmission electron microscope (Thermo Fisher Scientific) operating at 200 kV, using the EPU automated image acquisition software (Thermo Fisher Scientific). Movies were collected on a Falcon 4i direct electron detector operating in counting mode at a nominal magnification of 150,000× (0.95 Å/pixel) using defocus range of −1.0 to −2.75 µm. Movies were collected over a 4.3-s exposure and a total dose of ∼40 e-/Å^2^.

Single-particle cryo-EM data of the stabilized ectodomain were acquired on a Glacios transmission electron microscope (Thermo Fisher Scientific) operating at 200 kV, using the EPU automated image acquisition software (Thermo Fisher Scientific). Movies were collected on a Falcon 4i direct electron detector operating in counting mode at a nominal magnification of 150,000× (0.95 Å/pixel) using defocus range of −1.0 to −2.75 µm. Movies were collected over a 4.3-s exposure and a total dose of ∼40 e-/Å^2^. All movies were motion-corrected and dose-weighted with MotionCorr2 [61] and the aligned micrographs were used to estimate the defocus values with patchCTF within cryoSPARC [62]. CryoSPARC blob picker was used for automated particle picking, and the resulting particles were used to obtain initial 2D references. Five initial *ab initio* 3D models were generated in cryoSPARC and the one with the more particles was selected to perform a heterogeneous refinement into three classes without imposing any symmetry. The particles from the best class (which generated a map with more density) were subjected to a 3D classification into 5 classes using cryoSPARC without initial volumes or masks. The result with best density across all protomers was chosen for a non-uniform refinement with C3 symmetry in cryoSPARC [62]. The final map was sharpened with DeepEMhancer [63].

Model building started with the domains I, II and III (missing the HRA refolding region) from the pre-fusion model, and later from the Alphafold multimer model of the trimeric full-length Env. The individual domains were fitted into the sharpened EM map using UCSF Chimera [64]. The regions connecting the individual domains were deleted and manually re-built. This initial model was refined with one round of real space refinement including simulated annealing in Phenix [65] followed by iterative rounds of manual building and real-space and B-factor refinement in Coot [59] and Phenix, with secondary structure restraints. Symmetry operators were obtained from the EM map with the map-symmetry tool in Phenix, and they were used to place the three copies of the protomer within the trimeric map with the apply-NCS-operators program in Phenix. Validation of model coordinates was performed using MolProbity [60].

### Cells, sequences, and production of Foamy virus viral vectors

The four-component foamy virus vector (FVV) system (plasmids pcoPG, pcoPP, EnvGI-SUGII, and pcu2MD9-BGAL (a transfer plasmid encoding for β-galactosidase)) has been described [66]. The EnvGI-SUGII plasmid encodes a full-length gorilla Env that comprises the LP and TM sequences from the zoonotic SFVggo_huBAD468 (JQ867465) and the surface subunit (SU) from the SFVggo_huBAKK74 (JQ867464). We refer to this Env as WT. Mutation E630P was introduced in WT Env. FVVs were produced by co-transfection of four plasmids (gag:env:pol:transgene β-galactosidase) at a ratio of 7:2:3:26. Three micrograms total DNA and 8 μl polyethyleneimine (JetPEI, Polyplus, Ozyme) were added to 0.5 × 10^6^ HEK 293T cells seeded in 6-well plates. Supernatants were collected 48 h post transfection, clarified at 1500 × g for 10 min, and stored as single-use aliquots at −80 °C. Vector infectivity was determined by transducing BHK-21 cells with serial five-fold dilutions of vectors and detecting β-galactosidase expression after 72 h of culture at 37 °C. Plates were fixed with 0.5% glutaraldehyde in phosphate-buffered saline (PBS) for 10 min at room temperature, washed with PBS and stained with 150 μl X-gal solution containing 2mM MgCl_2_, 10mM potassium ferricyanide, 10mM potassium ferrocyanide and 0.8 mg/ml 5-bromo-4-chloro-3indolyl-B-D-galactopyranoside in PBS for 3 h at 37 °C. Counting was done on a S6 Ultimate Image UV analyzer (CTL Europe), with one blue cell defined as one infectious unit. Cell transduction by FVV is a surrogate for viral infectivity and FVV titers were expressed as infectious units/ml.

The yield of FVV particles was estimated by the quantification of particle-associated transgene RNA as described in [34]. FVV RNAs were extracted from raw cell supernatants with QIAamp Viral RNA Extraction Kit (Qiagen). RNAs were treated with DNA free kit (Life Technologies), retro-transcribed with Maxima H Minus Reverse Transcriptase (Thermo Fischer Scientific) using random primers (Thermo Fischer Scientific), according to manufacturer’s instructions. qPCR was performed on cDNA using BGAL primers (BGAL_F 5’ AAACTCGC AAGCCGACTGAT 3’ and BGAL_R 5’ ATATCGCGGCTCAGTTCGAG 3’) with a 10 min denaturation step at 95 °C and 40 amplification cycles (15 s at 95 °C, 20 s at 60 °C and 30 s at 72 °C) carried out with an Eppendorf realplex2 Mastercycler. A standard curve prepared with serial dilutions of pcu2MD9-BGAL plasmid was used to determine the copy number of FVVs. Results were expressed as vector particles/ml, considering that each particle carries 2 copies of the transgene.

The capacity of FVVs to bind to HT1080 cells was tested by incubating cells with FVV particles (1, 10, and 100 particles/cell) on ice for 1 h. Cells were washed three times with PBS to eliminate unbound FVVs and RNAs were extracted using RNeasy plus mini kit (Qiagen) according to manufacturer’s protocol. RT was performed as described for FVVs RNA quantification. Bound FVV were quantified by qPCR of *bgal* gene as described for vector titration; cells were quantified by a qPCR amplifying the *hgapdh* gene with the following primers: hGAPDH_F 5’ GGAGCGAGATCCCTCCAAAAT 3’ and hGAPDH_R 5’ GGCTGTTGTCATACTTCTCATGG 3’. The qPCR reaction conditions were the same as those used to amplify the *bgal* gene. Relative mRNA expression of *bgal* versus *hgapdh* was calculated using the –ΔΔCt method, and relative binding as 2–ΔΔCt.

### Alphafold multimer prediction of full-length FV Env trimer

For the generation of the FV Env model we used ColabFold v1.5.1 [67] with the weights of AlphaFold multimer v3 [35]. The multiple sequence alignment was generated using the mmseqs2 [68] and the uniref30 database [69], resulting in a multiple sequence alignment (MSA) of 79 sequences. The template search did not return any matches, so the model was generated from only MSAs as an input. All the models were predicted using 20 recycles on an NVIDIA V100 GPU (taking about 5 hours per prediction).

## Statistics

The infectious titers, particle concentration, percentages of infectious particles and quantity of bound FVVs carrying WT and E630P Env variant were compared using two-way paired t-test, with p-values indicated on the graph.

## Data availability

The coordinates for the cryo-EM structural models have been deposited to the RCSB protein databank under PDB accession codes 8RM0 (pre-fusion Env) and 8RM1 (post-fusion Env). The EM maps have been deposited to the EMDB under accession codes EMD-19347 (pre-fusion Env) and EMD-19348 (post-fusion Env). The Alphafold multimer model of FV Env trimer in the pre-fusion state is available in ModelArchive (modelarchive.org) with the accession code ma-q5jmc.

## Acknowledgements

This project was funded by the Integrative Biology of Emerging Infectious Diseases Laboratory of Excellence [ANR-10-LABX62-IBEID, Intra-Labex Grant, M.B] and the ‘Programme de recherche transversal from Institut Pasteur’ [PTR2020-353 ZOOFOAMENV, Florence Buseyne]. The funding agencies had no role in the study design, generation of results, or writing of the manuscript. We thank the staff from the Nanocore Imagining Facility at Institut Pasteur for technical assistance during EM data collection. We are grateful to Jan Hellert and Pablo Guardado Calvo for the discussions and advice, and for critical reading of the manuscript.

I dedicate this manuscript to the memory of Robert Lamb, for his advice along the years and for what I had learned about paramyxoviruses while I was a postdoc in the Jardetzky lab (M.B.).

## Author contributions

Conceptualization: IF, MB, Florence Buseyne

Realization of the assays: IF, François Bontems, YC, DB

Resources: MB, Florence Buseyne, FAR, AG

Supervision: MB, Florence Buseyne

Writing—original draft: IF

Writing—review & editing: MB, FAR, IF, Florence Buseyne, François Bontems, CG, BC

Funding acquisition: MB, Florence Buseyne

