## Supplementary Info for "Structures of the Foamy virus fusion protein reveal an unexpected link with the F protein of paramyxo- and pneumoviruses"

### Supplementary information

#### Figure S1: Post-fusion Env expression and structural features

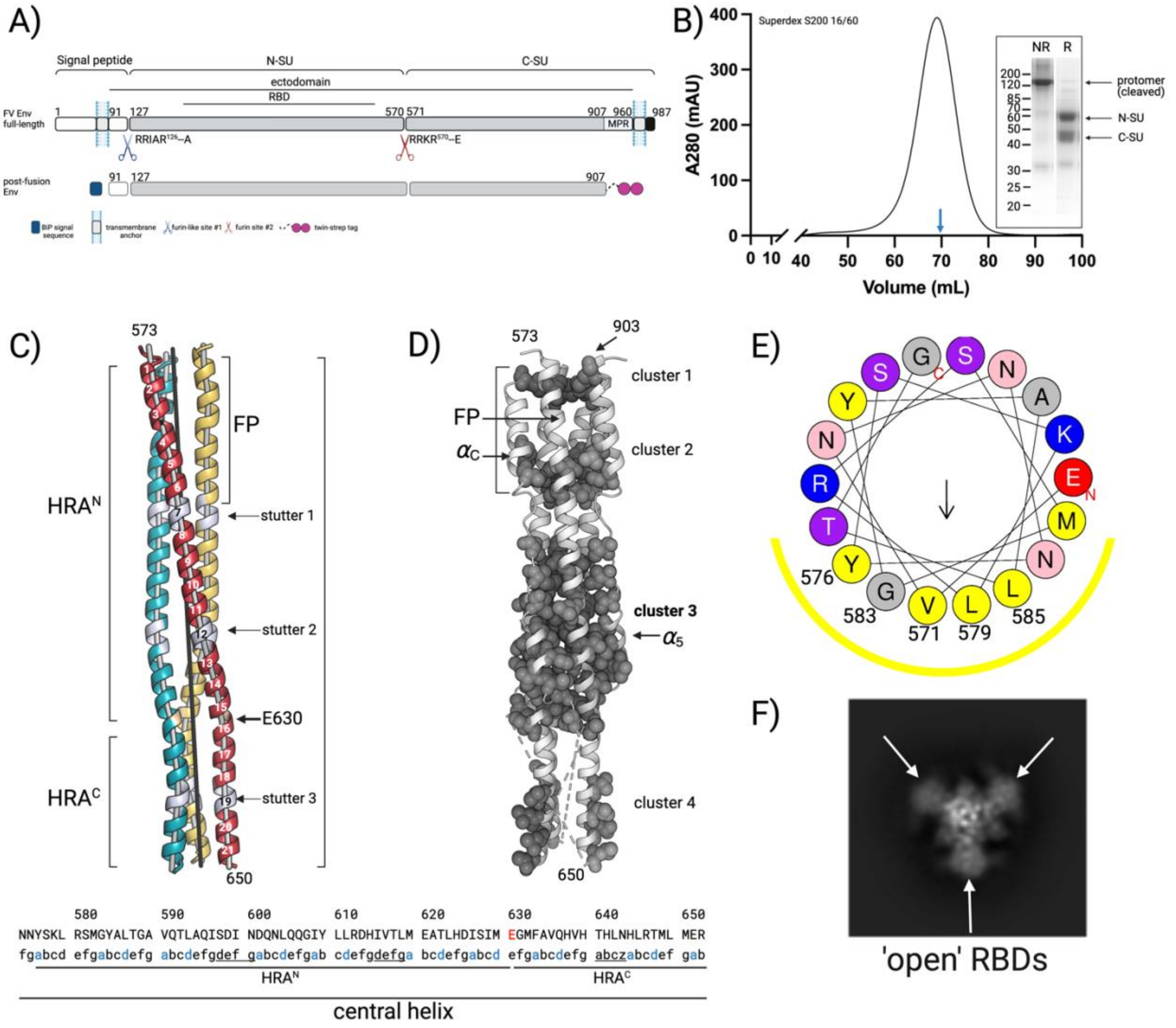

**A)** Schematic representation of the full-length FV Env and the expression construct used for determination of the post-fusion Env ectodomain structure.

**B)** Size exclusion chromatogram of the affinity purified post-fusion Env ectodomain. The fraction from the middle of the peak (labeled with blue arrow) was analyzed by SDS-PAGE under non-reducing (NR) and reducing (R) conditions.

**C)** The trimeric coiled coil formed by the central helices, and composed of HRA$^N$ and HRA$^C$ segments, is shown with each chain colored differently. The turns are labeled with numbers on the dark red protomer. The axes of the $\alpha$ helices and the coiled coil are represented as grey and black sticks, respectively. The residues corresponding to the *a* and *d* positions in a coiled coil and identified by program Twister [1] are highlighted below as blue letters. The residues forming 3 stutters, which break the HR pattern, are underlined, and indicated on the structure above.

**D)** The trimeric coiled coil formed by the central helices is shown as grey cartoon. The residues forming hydrophobic clusters are shown in space fill model.

**E)** Helical wheel projections for the amino acid residues corresponding to the region containing the FP (residues Glu 570 – Gly 588) are shown to illustrate its amphipathic nature. The non-polar side of the helix is indicated with yellow line.

**F)** A 2D class showing the RBDs (white arrows) in an 'open' rearrangement.

#### Figure S2: Cryo-EM SPA and generation of map for the post-fusion Env structure

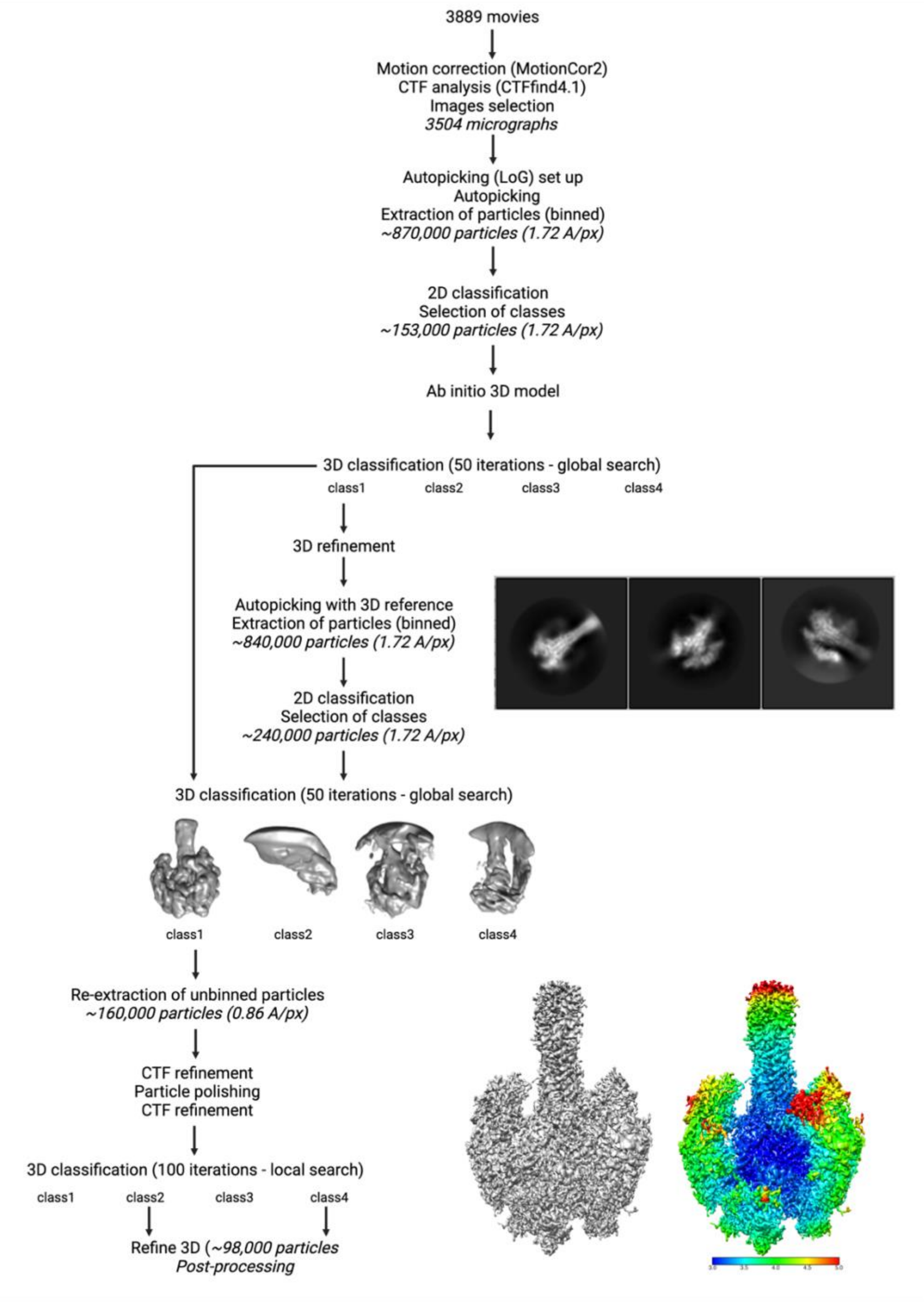

Scheme illustrating the steps followed to process the collected cryo-EM data of the FV Env WT ectodomain. Selected 2D class averages, the final map, and a local resolution graphic (gradient blue to red) are also shown.

#### Figure S3: Structure and secondary structure topology of domains I, II and III of post-and pre-fusion FV Env

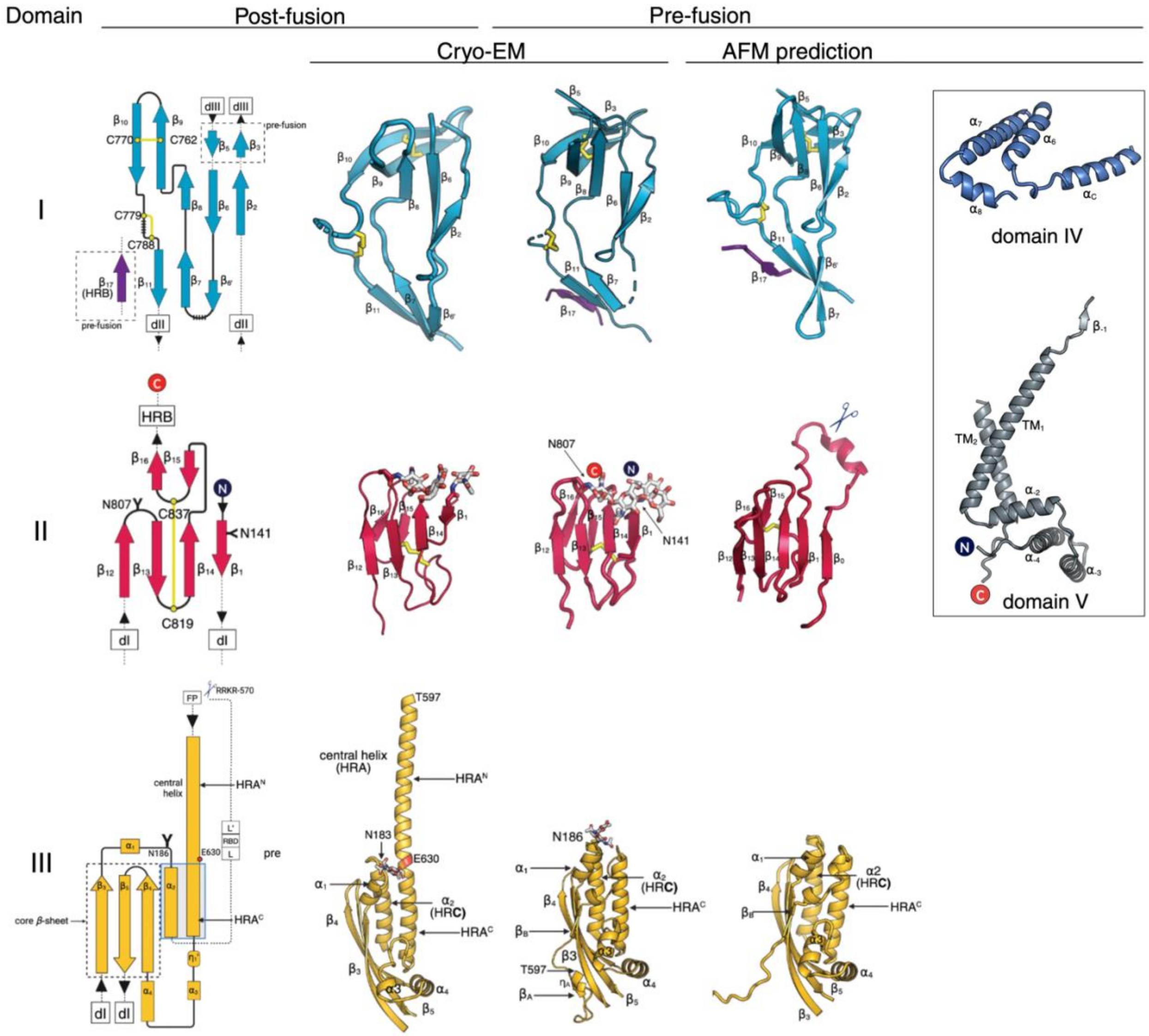

Secondary structure topology diagrams and cartoon representations of domains I, II and III are shown for the post- and pre-fusion FV Env ectodomain structure obtained by cryo-EM. The cartoon representations for domains I to V of the Alphafold multimer (AFM) predicted model are shown on the right side of the panel for comparison. The color scheme corresponds to the one shown on Fig. 1C. Domain boundaries are defined as: domain I (residues 145-161 and 713-796), domain II (residues 139-144 and 797-844), domain III (residues 162-206 and 597-712), domain IV (residues 886-956), and domain V (residues 1-99 and 957-987). Of note, the $\beta$ sheet of domain I is augmented, in the pre-fusion state, by strand $\beta_{17}$ from HRB and few residues of long strands $\beta_3$ and $\beta_5$ that belong to domain III (see Fig. S6). According to the AFM model, the $\alpha$-helices forming membrane spanning segments are: residues 66-100 ($TM_1$) and residues 957-979 ($TM_2$). The FP and HRB are not included because they interact with different domains in the pre- and post-fusion conformations.

#### Figure S4: Pre-fusion Env construct design and characterization.

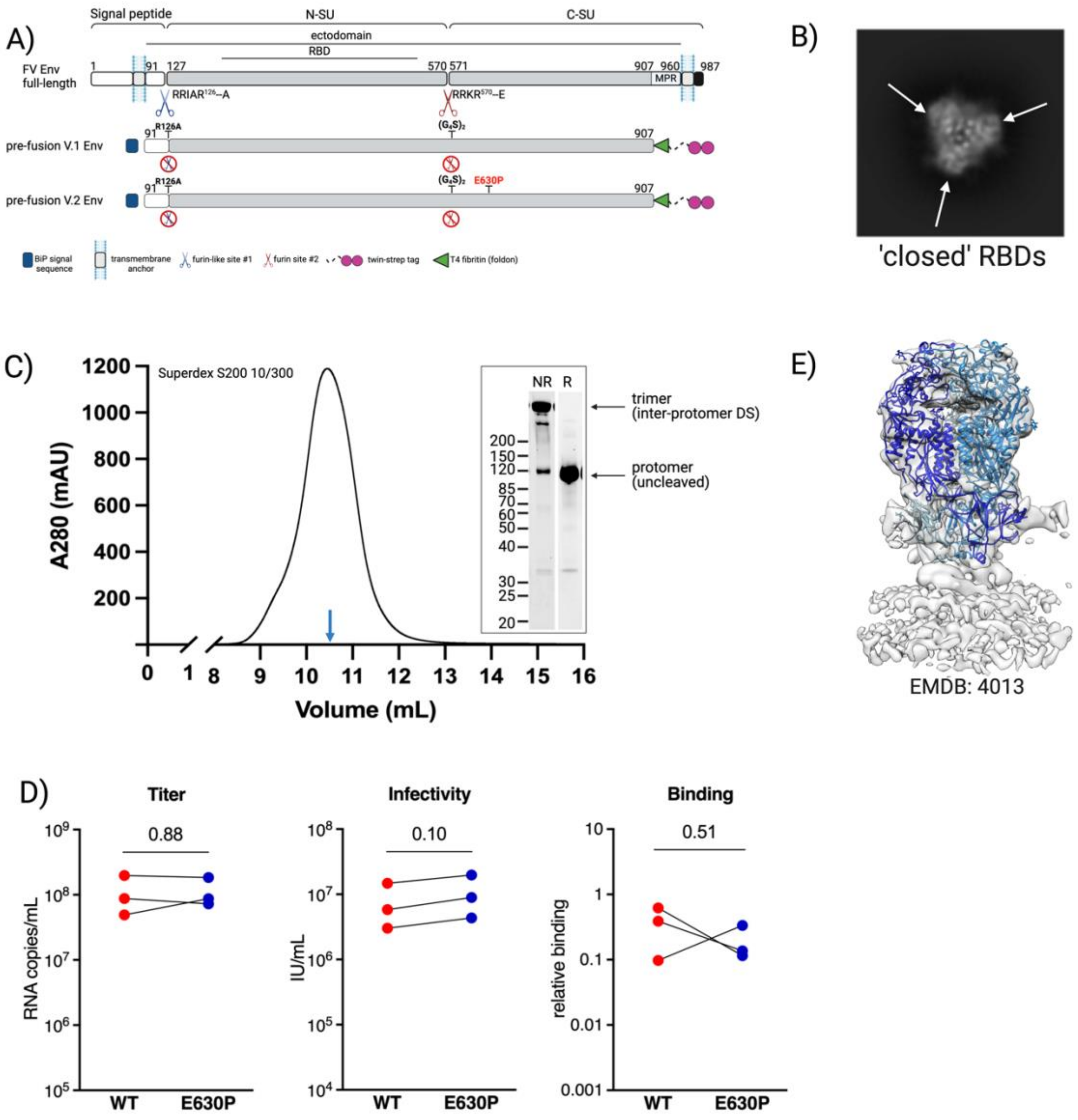

**A)** Schematic representation of the full-length FV Env and the expression constructs V.1 and V.2 used for determination of the pre-fusion Env ectodomain structure (see Methods).

**B)** A 2D class calculated for the pre-fusion ectodomain, showing the RBDs (white arrows) in a 'closed' rearrangement.

**C)** Size exclusion chromatogram of the affinity purified pre-fusion Env ectodomain V.2. The fraction from the middle of the peak (labeled with blue arrow) was analyzed by SDS-PAGE under non-reducing (NR) and reducing (R) conditions.

**D)** The titer, infectivity, and binding of E630P Env viral vector particles is shown. Three batches of FVVs carrying WT or E630P Envs were produced. Titers of FVV particles in supernatants were quantified by RT-qPCR. Infectivity was quantified through transduction of susceptible cells. Vector particles binding to HT1080 cells was quantified by incubating particles with cells on ice for 1 h, before washing and quantifying the remaining vector transgene and cellular hgapdh by RT-qPCR; results are shown for the 10 particle/cell dose and are expressed as relative binding. Each batch is represented with a single dot (red and blue for vectors carrying WT and E630P Envs, respectively). The FVVs carrying the WT and E630P Envs were compared using the two-way paired t-test, with p-values indicated on the graph.

**E)** Fitting of our pre-fusion cryo-EM structure in the map obtained for the full length Env from viral vectors [2]. The protomers are highlighted with 3 shades of blue.

#### Figure S5: Cryo-EM SPA and generation of map for the pre-fusion Env structure

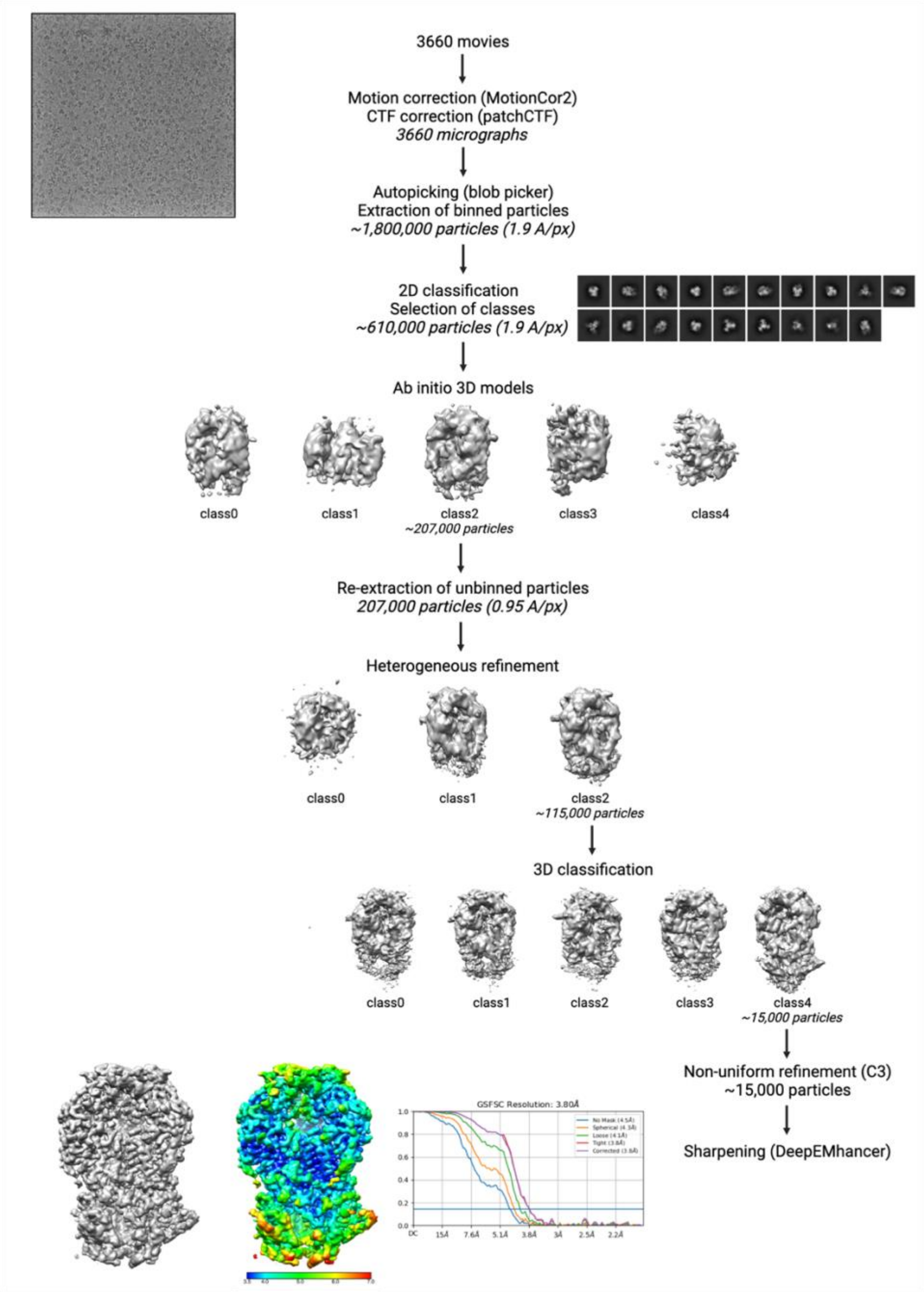

Scheme detailing the steps followed to process the cryo-EM data collected for the stabilized FV Env ectodomain V.2 (Fig. S4A). A micrograph with particles, selected 2D class averages, the final sharpened map, a local resolution graphic, and the GSFSC (gold standard Fourier shell correlation) resolution plot are shown.

#### Figure S6: Sequence alignment of FV Env

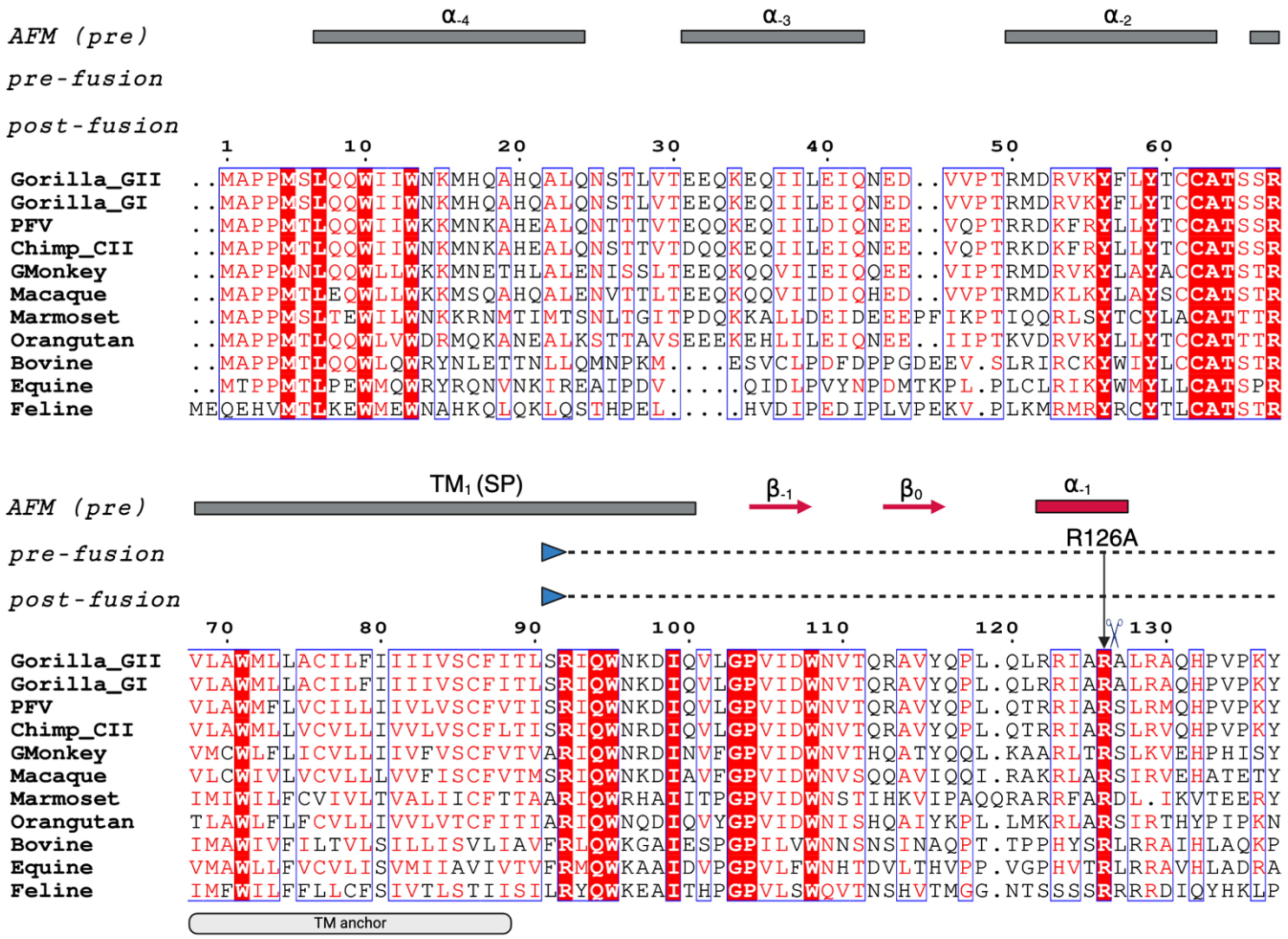

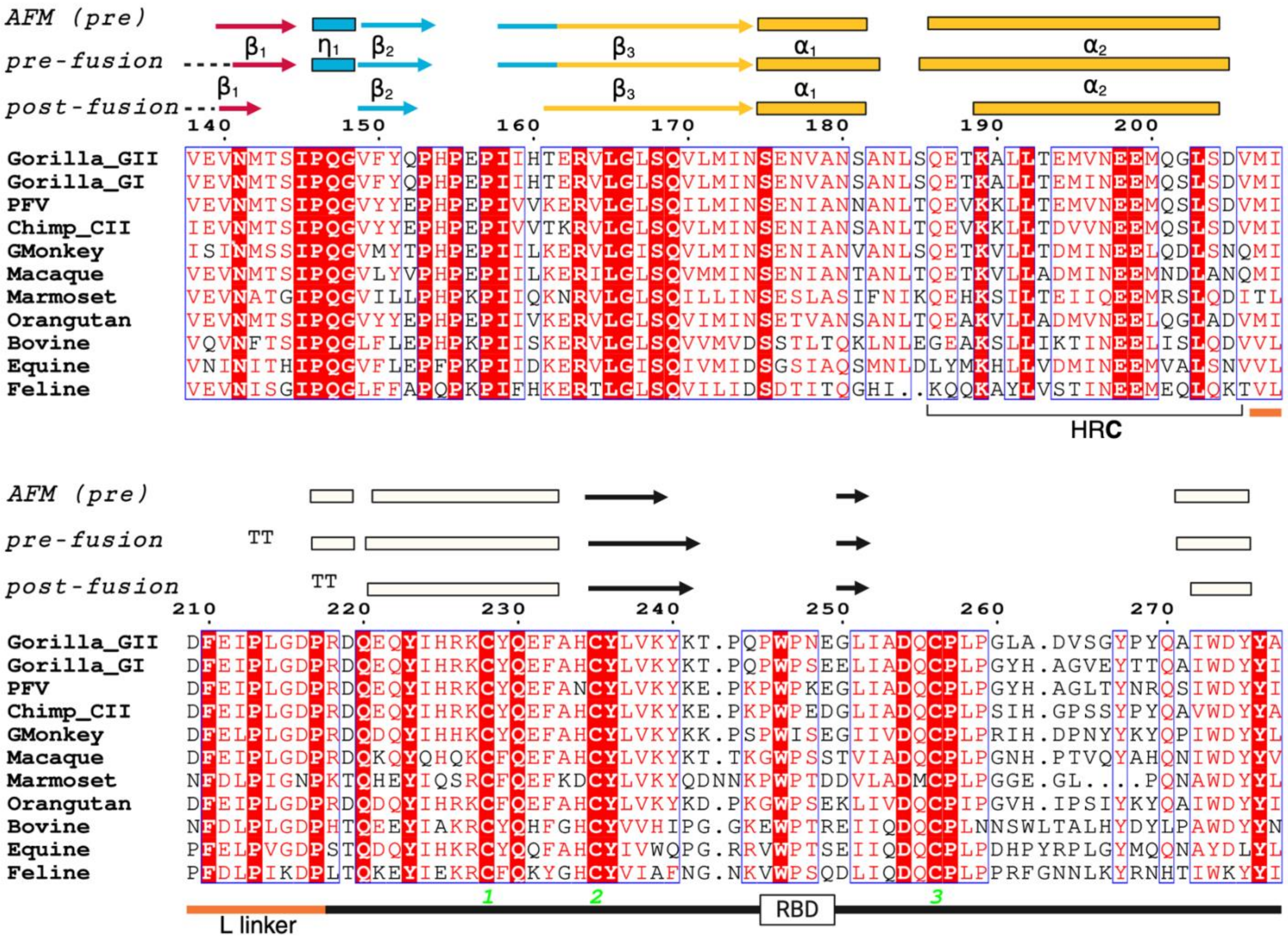
AFM (pre)
pre-fusion
post-fusion
β1
η1
β2
β3
α1
α2
HRC
TT
Gorilla_GII
Gorilla_GI
PFV
Chimp_CII
GMonkey
Macaque
Marmoset
Orangutan
Bovine
Equine
Feline
L linker
RBD

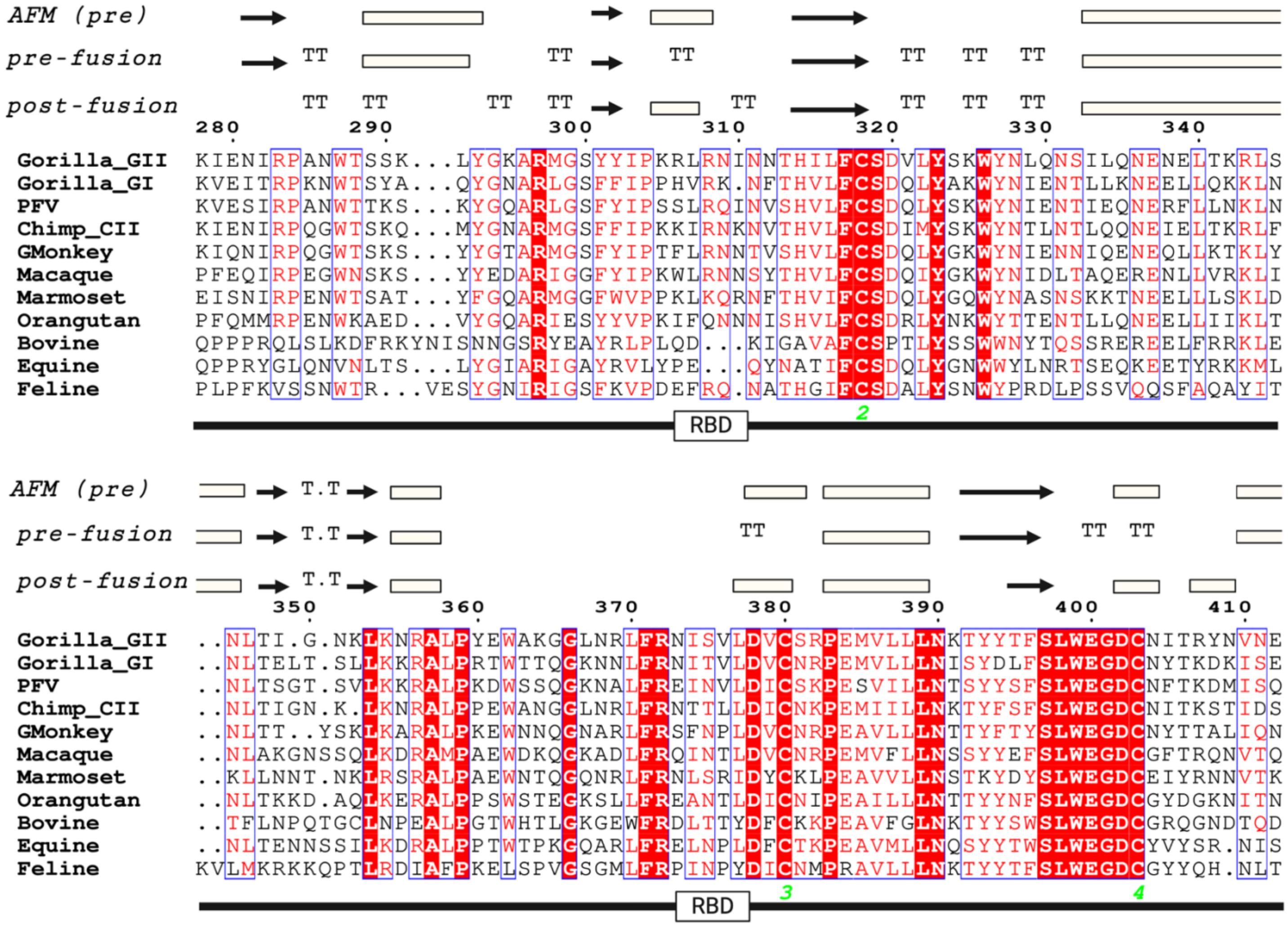
AFM (pre)
pre-fusion
post-fusion
TT
280
290
300
310
320
330
340
Gorilla_GII KIENIRPANWTSSK...LYGKARMGSYYIPKRLRNINNTHILFCSDVLYSKWYNLQNSILQNENELTKRLS
Gorilla_GI KVEITRPKNWTSYA...QYGNARLGSFFIPPHVRK.NFTHVLFCSDQLYAKWYNIENTLLKNEELLQKKLN
PFV KVESIRPANWTTKS...KYGQARLGSFYIPSSLRQINVSHVLFCSDQLYSKWYNIENTIEQNERFLLNKLN
Chimp_CII KIENIRPQGWTSKQ...MYGNARMGSFFIPKKIRNKNVTHVLFCSDIMYSKWYNTLNTLQQNEIELTKRLF
GMonkey KIQNIRPQGWTSKS...YYGTARMGSFYIPTFLRNNTVSHVLFCSDQLYGKWYNIENNIQENEQLLKTKLY
Macaque PFEQIRPEGWNSKS...YYEDARIGGFYIPKWLRNNSYTHVLFCSDQLYGKWYNIDLTAQERENLLVRKLI
Marmoset EISNIRPENWTSAT...YFGQARMGGFWVPPKLKQRNFTHVIFCSDQIYGQWYNASNSKKTNEELLLSKLD
Orangutan PFQMMRPENWKAED...VYGQARIESYYVPKIFQNNNISHVLFCSDRLYNKWYTTENTLLQNEELLIIKLT
Bovine QPPPRQLSLKDFRKYNISNNGSRYEAYRLPLQD...KIGAVAFCSPTLYSSWWNYTQSSREREELFRRKLE
Equine QPPPRYGLQNVNLTS...LYGIARIGAYRVLYPE...QYNATIFCSDQLYGNWWYLNRTSEQKEETYRKKML
Feline PLPFKVSSNWTR...VESYGNIRIGSFKVPDEFRQ.NATHGIFCSDALYSNWYPRDLPSSVQQSFAQAYIT
2
RBD
AFM (pre)
pre-fusion
post-fusion
T.T
TT
350
360
370
380
390
400
410
Gorilla_GII ..NLTI.G.NKLKNRALPYEWAKGGLNRLFRNISVLDVCSRPEMVLLLNKTYYTFSLWEGDCNITRYNVNE
Gorilla_GI ..NLTELT.SLLKKRALPRTWTTQGKNNLFRNITVLDVCNRPEMVLLLNISYDLFSLWEGDCNYTKDKISE
PFV ..NLTSGT.SVLKKRALPKDWSSQGKNALFREINVLDICSKPESVILLNTSYYSFSLWEGDCNFTKDMISQ
Chimp_CII ..NLTIGN.K.LKNRALPPEWANGGLNRLFRNTTLLDICNKPEMIILLNKTYFSFSLWEGDCNITKSTIDS
GMonkey ..NLTT..YSKLKARALPKEWNNQGNARLFRSFNPLDVCNRPEAVLLLNTTYFTYSLWEGDCNYTTALIQN
Macaque ..NLAKGNSSQLKDRAMPAEWDKQGKADLFRQINTLDVCNRPEMVFLLNSSYYEFSLWEGDCGFTRQNVTQ
Marmoset ..KLLNNT.NKLRSRALPAEWNTQGQNRLFRNLSRIDYCKLPEAVVLLNSTKYDYSLWEGDCEIYRNNVTK
Orangutan ..NLTKKD.AQLKERALPPSWSTEGKSLLFREANTLDICNIPEAILLLNTTYYNFSLWEGDCGYDGKNITN
Bovine ..TFLNPQTGCLNPEALPGTWHTLGKGEWFRDLTTYDFCKKPEAVFGLNKTYYSWSLWEGDCGRQGNDTQD
Equine ..NLTENNSSILKDRALPPTWTPKGQARLFRELNPLDFCTKPEAVMLLNQSYYTWSLWEGDCYVYSR.NIS
Feline KVLMKRKKQPTLRDIAFPKELSPVGSGMLFRPINPYDICNMPRAVLLLNKTYYTFSLWEGDCGYYQH.NLT
3
4
RBD

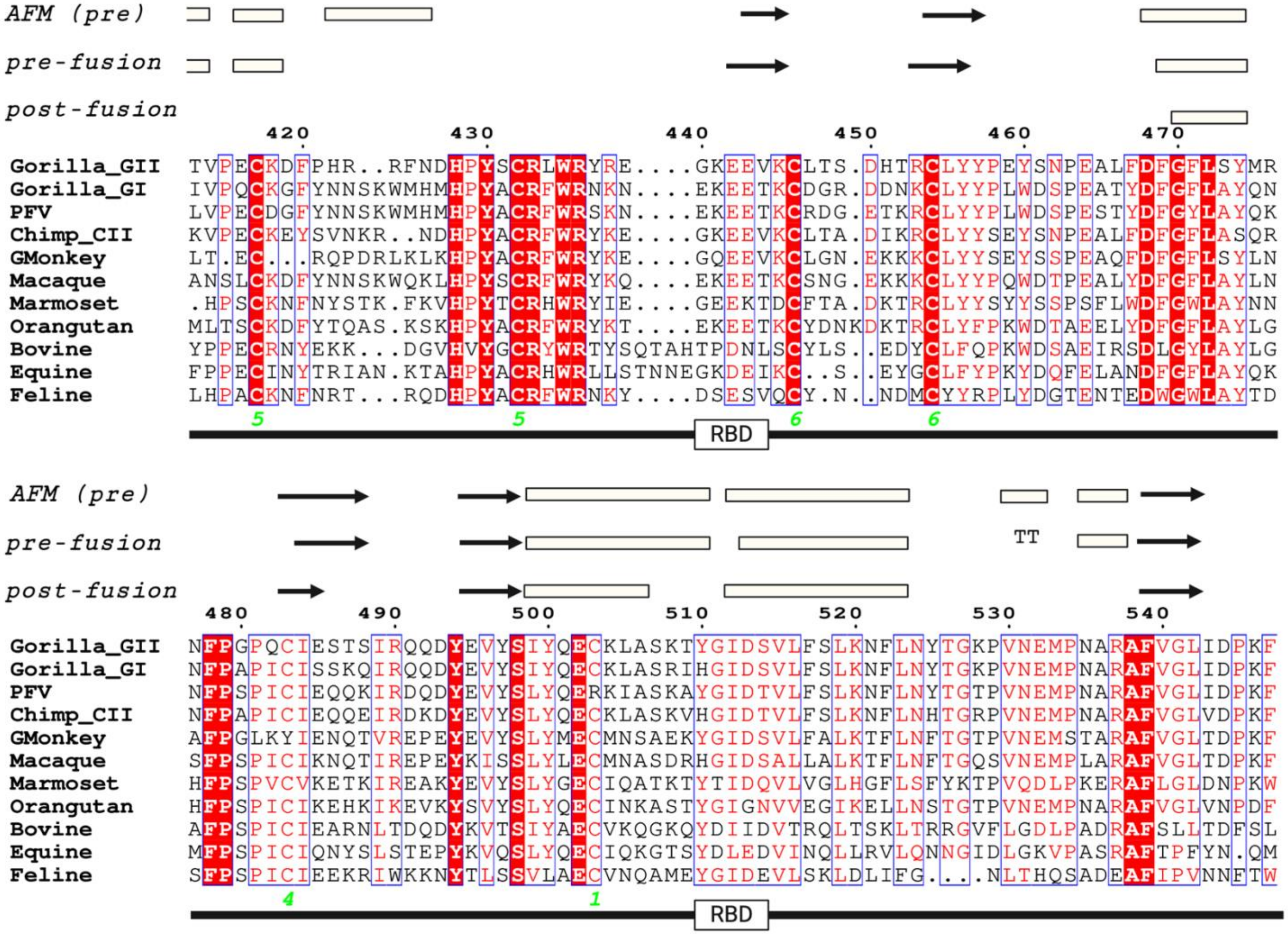
AFM (pre)
pre-fusion
post-fusion
420 430 440 450 460 470
Gorilla_GII TVPECKDFPHR..RFNDHPYSCRLWRYRE....GKEEVKCLTS.DHTRCLYYPEYSNPEALFDFGFLSYMR
Gorilla_GI IVPQCKGFYNNSKWMHMHPYACRFWRNKN....EKEETKCDGR.DDNKCLYYPLWDSPEATYDFGFLAYQN
PFV LVPECDGFYNNSKWMHMHPYACRFWRSKN....EKEETKCRDG.ETKRCLYYPLWDSPESTYDFGYLAYQK
Chimp_CII KVPECKEYSVNKR..NDHPYACRFWRYKE....GKEEVKCLTA.DIKRCLYYSEYSNPEALFDFGFLASQR
GMonkey LT.EC...RQPDRLKLKHPYACRFWRYKE....GQEEVKCLGN.EKKKCLYYSEYSSPEAQFDFGFLSYLN
Macaque ANSLCKDFYNNSKWQKLHPYSCRFWRYKQ....EKEETKCSNG.EKKKCLYYPQWDTPEALYDFGFLAYLN
Marmoset .HPSCKNFNYSTK.FKVHPYTCRHWRYIE....GEEKTDCFTA.DKTRCLYYSYYSSPSFLWDFGWLAYNN
Orangutan MLTSCKDFYTQAS.KSKHPYACRFWRYKT....EKEETKCYDNKDKTRCLYFPKWDTAEELYDFGFLAYLG
Bovine YPPECRNYEKK...DGVHVYGCRYWRTYSQTAHTPDNLSCYLS..EDYCLFQPKWDSAEIRSDLGYLAYLG
Equine FPPECINYTRIAN.KTAHPYACRHWRLLSTNNEGKDEIKC..S..EYGCLFYPKYDQFELANDFGFLAYQK
Feline LHPACKNFNRT...RQDHPYACRFWRNKY....DSESVQCY.N..NDMCYYRPLYDGTENTEDWGWLAYTD
5 5 6 6
RBD
AFM (pre)
pre-fusion
TT
post-fusion
480 490 500 510 520 530 540
Gorilla_GII NFPGPQCIESTSIRQQDYEVYSIYQECKLASKTYGIDSVLFSLKNFLNYTGKPVNEMPNARAFVGLIDPKF
Gorilla_GI NFPAPICISSKQIRQQDYEVYSIYQECKLASRIHGIDSVLFSLKNFLNYTGKPVNEMPNARAFVGLIDPKF
PFV NFPSPICIEQQKIRDQDYEVYSLYQERKIASKAYGIDTVLFSLKNFLNYTGTPVNEMPNARAFVGLIDPKF
Chimp_CII NFPAPICIEQQEIRDKDYEVYSLYQECKLASKVHGIDTVLFSLKNFLNHTGRPVNEMPNARAFVGLVDPKF
GMonkey AFPGLKYIENQTVREPEYEVYSLYMECMNSAEKYGIDSVLFALKTFLNFTGTPVNEMSTARAFVGLTDPKF
Macaque SFPSPICIKNQTIREPEYKISSLYLECMNASDRHGIDSALLALKTFLNFTGQSVNEMPLARAFVGLTDPKF
Marmoset HFPSPVCVKETKIREAKYEVYSLYGECIQATKTYTIDQVLVGLHGFLSFYKTPVQDLPKERAFLGLDNPKW
Orangutan HFPSPICIKEHKIKEVKYSVYSLYQECINKASTYGIGNVVEGIKELLNSTGTPVNEMPNARAFVGLVNPDF
Bovine AFPSPICIEARNLTDQDYKVTSIYAECVKQGKQYDIIDVTRQLTSKLTRRGVFLGDLPADRAFSLLTDFSL
Equine MFPSPICIQNYSLSTEPYKVQSLYQECIQKGTSYDLEDVINQLLRVLQNNGIDLGKVPASRAFTPFYN.QM
Feline SFPSPICIEEKRIWKKNYTLSSVLAECVNQAMEYGIDEVLSKLDLIFG...NLTHQSADEAFIPVNNFTW
4 1
RBD

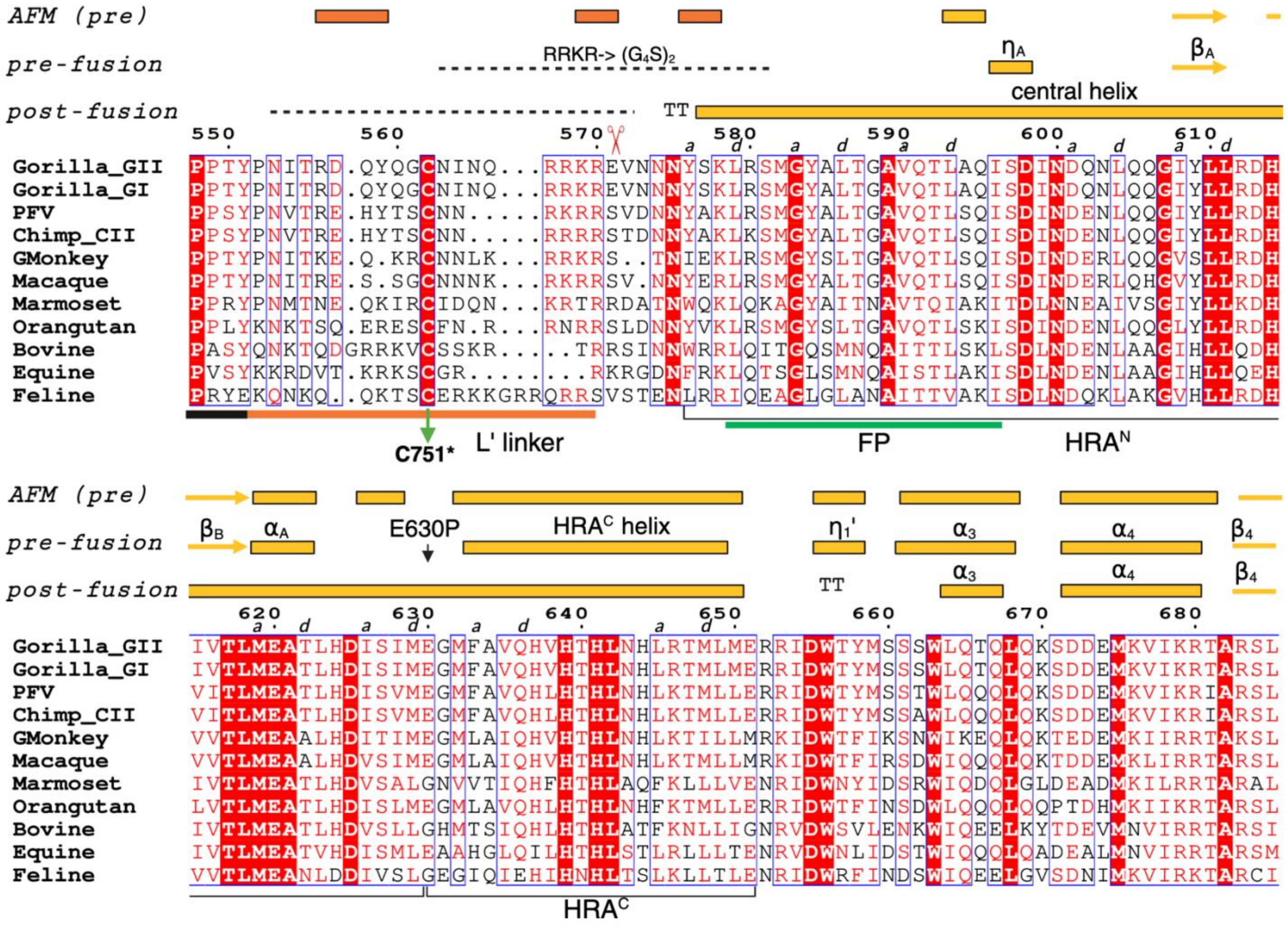
AFM (pre)
pre-fusion
post-fusion
RRKR-> (G4S)2
ηA
βA
central helix
TT
550
560
570
580
590
600
610
Gorilla_GII PPTYPNITRD.QYQGCNINQ...RRKREVNNNYSKLRSMGYALTGAVQTLAQISDINDQNLQQGIYLLRDH
Gorilla_GI PPTYPNITRD.QYQGCNINQ...RRKREVNNNYSKLRSMGYALTGAVQTLAQISDINDQNLQQGIYLLRDH
PFV PPSYPNVTRE.HYTSCNN.....RKRRSVDNNYAKLRSMGYALTGAVQTLSQISDINDENLQQGIYLLRDH
Chimp_CII PPSYPNVTRE.HYTSCNN.....RRRRSTDNNYAKLKSMGYALTGAVQTLSQISDINDENLQQGIYLLRDH
GMonkey PPTYPNITKE.Q.KRCNNLK...RRKRS..TNIEKLRSMGYSLTGAVQTLSQISDINDERLQQGVSLLRDH
Macaque PPTYPNITRE.S.SGCNNNK...RKRRSV.NNYERLRSMGYALTGAVQTLSQISDINDERLQHGVYLLRDH
Marmoset PPRYPNMTNE.QKIRCIDQN...KRTRRDATNWQKLQKAGYAITNAVTQIAKITDLNNEAIVSGIYLLKDH
Orangutan PPLYKNKTSQ.ERESCFN.R...RNRRSLDNNYVKLRSMGYSLTGAVQTLSKISDINDENLQQGLYLLRDH
Bovine PASYQNKTQDGRRKVCSSKR.....TRRSINNWRRLQITGQSMNQAITTLSKLSDLNDENLAAGIHLLQDH
Equine PVSYKKRDVT.KRKSCGR........RKRGDNFRKLQTSGLSMNQAISTLAKISDLNDENLAAGIHLLQEH
Feline PRYEKQNKQ..QKTSCERKKGRRQRRSVSTENLRRIQEAGLGLANAITTVAKISDLNDQKLAKGVHLLRDH
C751*
L' linker
FP
HRA^N
AFM (pre)
pre-fusion
post-fusion
βB
αA
E630P
HRA^C helix
η1'
α3
α4
β4
TT
620
630
640
650
660
670
680
Gorilla_GII IVTLMEATLHDISIMEGMFAVQHVHTHLNHLRTMLMERRIDWTYMSSSWLQTQLQKSDDEMKVIKRTARSL
Gorilla_GI IVTLMEATLHDISIMEGMFAVQHVHTHLNHLRTMLMERRIDWTYMSSSWLQTQLQKSDDEMKVIKRTARSL
PFV VITLMEATLHDISVMEGMFAVQHLHTHLNHLKTMLLERRIDWTYMSSTWLQQQLQKSDDEMKVIKRIARSL
Chimp_CII VITLMEATLHDISVMEGMFAVQHLHTHLNHLKTMLLERRIDWTYMSSAWLQQQLQKSDDEMKVIKRIARSL
GMonkey VVTLMEAALHDITIMEGMLAIQHVHTHLNHLKTILLMRKIDWTFIKSNWIKEQLQKTEDEMKIIRRTAKSL
Macaque VVTLMEAALHDVSIMEGMLAIQHVHTHLNHLKTMLLMRKIDWTFIRSDWIKEQLQKTDDEMKLIRRTARSL
Marmoset IVTLMEATLHDVSALGNVVTIQHFHTHLAQFKLLVENRIDWNYIDSRWIQDQLGLDEADMKILRRTARAL
Orangutan LVTLMEATLHDISLMEGMLAVQHLHTHLNHFKTMLLERRIDWTFINSDWLQQQLQQPTDHMKIIKRTARSL
Bovine IVTLMEATLHDVSLLGHMTSIQHLHTHLATFKNLLIGNRVDWSVLENKWIQEELKYTDEVMNVIRRTARSI
Equine IVTLMEATVHDISMLEAAHGLQILHTHLSTLRLLLTENRVDWNLIDSTWIQQQLQADEALMNVIRRTARSM
Feline VVTLMEANLDDIVSLGEGIQIEHIHNHLTSLKLLTLENRIDWRFINDSWIQEELGVSDNIMKVIRKTARCI
HRA^C

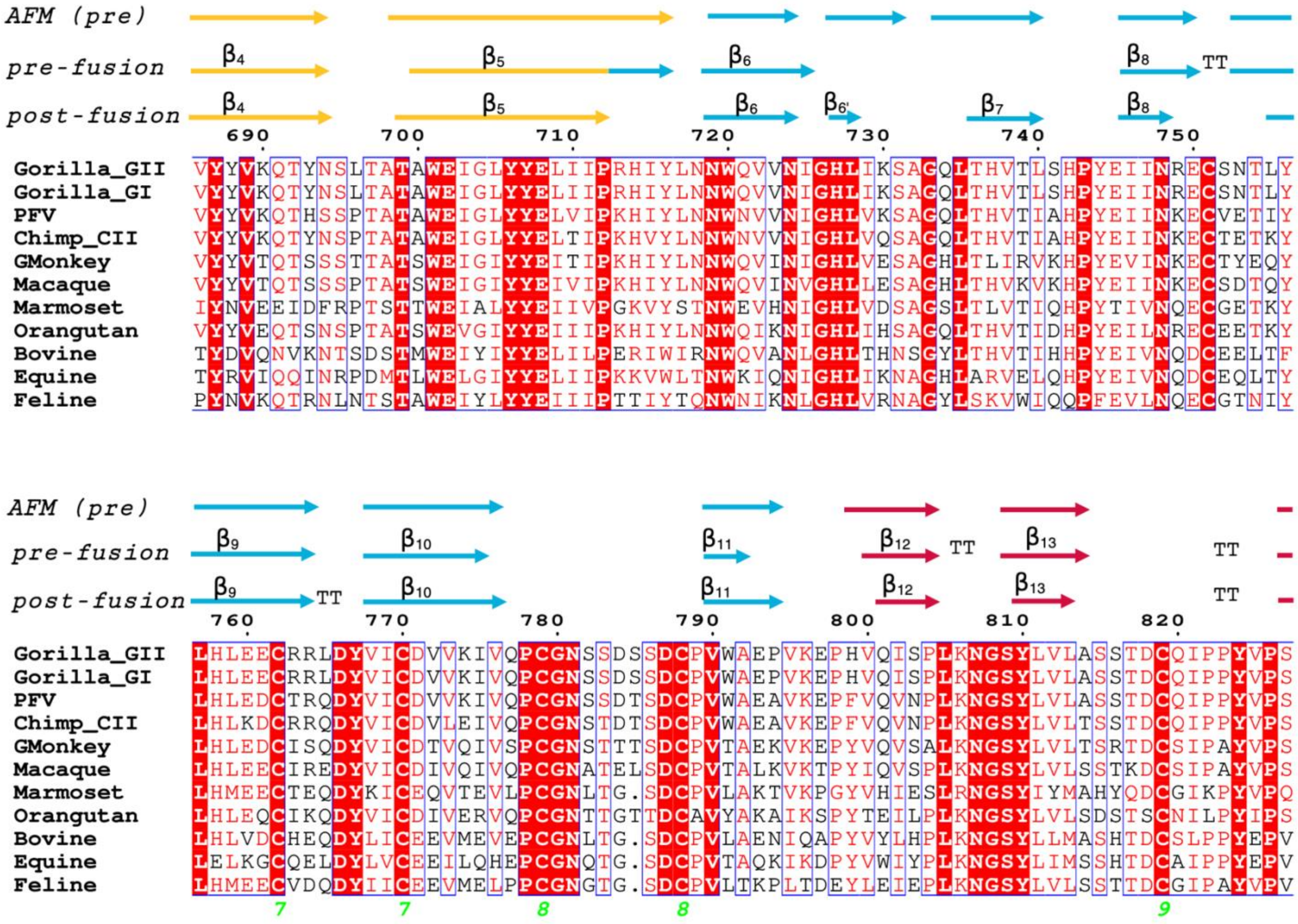
AFM (pre)
pre-fusion
post-fusion
β4
β5
β6
β6'
β7
β8
TT
690
700
710
720
730
740
750
Gorilla_GII VYYVKQTYNSLTATAWEIGLYYELIIPRHIYLNNWQVVNIGHLIKSAGQLTHVTLSHPYEIINRECSNTLY
Gorilla_GI VYYVKQTYNSLTATAWEIGLYYELIIPRHIYLNNWQVVNIGHLIKSAGQLTHVTLSHPYEIINRECSNTLY
PFV VYYVKQTHSSPTATAWEIGLYYELVIPKHIYLNNWNVVNIGHLVKSAGQLTHVTIAHPYEIINKECVETIY
Chimp_CII VYYVKQTYNSPTATAWEIGLYYELTIPKHVYLNNWNVVNIGHLVQSAGQLTHVTIAHPYEIINKECTETKY
GMonkey VYYVTQTSSSTTATSWEIGIYYEITIPKHIYLNNWQVINIGHLVESAGHLTLIRVKHPYEVINKECTYEQY
Macaque VYYVTQTSSSPTATSWEIGIYYEIVIPKHIYLNNWQVINVGHLLESAGHLTHVKVKHPYEIINKECSDTQY
Marmoset IYNVEEIDFRPTSTTWEIALYYEIIVPGKVYSTNWEVHNIGHLVDSAGSLTLVTIQHPYTIVNQECGETKY
Orangutan VYYVEQTSNSPTATSWEVGIYYEIIIPKHIYLNNWQIKNIGHLIHSAGQLTHVTIDHPYEILNRECEETKY
Bovine TYDVQNVKNTSDSTMWEIYIYYELILPERIWIRNWQVANLGHLTHNSGYLTHVTIHHPYEIVNQDCEELTF
Equine TYRVIQQINRPDMTLWELGIYYELIIPKKVWLTNWKIQNIGHLIKNAGHLARVELQHPYEIVNQDCEQLTY
Feline PYNVKQTRNLNTSTAWEIYLYYEIIIPTTIYTQNWNIKNLGHLVRNAGYLSKVWIQQPFEVLNQECGTNIY
AFM (pre)
pre-fusion
post-fusion
β9
β10
β11
β12
β13
TT
760
770
780
790
800
810
820
Gorilla_GII LHLEECRRLDYVICDVVKIVQPCGNSSDSSDCPVWAEPVKEPHVQISPLKNGSYLVLASSTDCQIPPYVPS
Gorilla_GI LHLEECRRLDYVICDVVKIVQPCGNSSDSSDCPVWAEPVKEPHVQISPLKNGSYLVLASSTDCQIPPYVPS
PFV LHLEDCTRQDYVICDVVKIVQPCGNSSDTSDCPVWAEAVKEPFVQVNPLKNGSYLVLASSTDCQIPPYVPS
Chimp_CII LHLKDCRRQDYVICDVLEIVQPCGNSTDTSDCPVWAEAVKEPFVQVNPLKNGSYLVLTSSTDCQIPPYVPS
GMonkey LHLEDCISQDYVICDTVQIVSPCGNSTTTSDCPVTAEKVKEPYVQVSALKNGSYLVLTSRTDCSIPAYVPS
Macaque LHLEECIREDYVICDIVQIVQPCGNATELSDCPVTALKVKTPYIQVSPLKNGSYLVLSSTKDCSIPAYVPS
Marmoset LHMEECTEQDYKICEQVTEVLPCGNLTG.SDCPVLAKTVKPGYVHIESLRNGSYIYMAHYQDCGIKPYVPQ
Orangutan LHLEQCIKQDYVICDIVERVQPCGNTTGTTDCAVYAKAIKSPYTEILPLKNGSYLVLSDSTSCNILPYIPS
Bovine LHLVDCHEQDYLICEEVMEVEPCGNLTG.SDCPVLAENIQAPYVYLHPLKNGSYLLMASHTDCSLPPYEPV
Equine LELKGCQELDYLVCEEILQHEPCGNQTG.SDCPVTAQKIKDPYVWIYPLKNGSYLIMSSHTDCAIPPYEPV
Feline LHMEECVDQDYIICEEVMELPPCGNGTG.SDCPVLTKPLTDEYLEIEPLKNGSYLVLSSTTDCGIPAYVPV
7
7
8
8
9

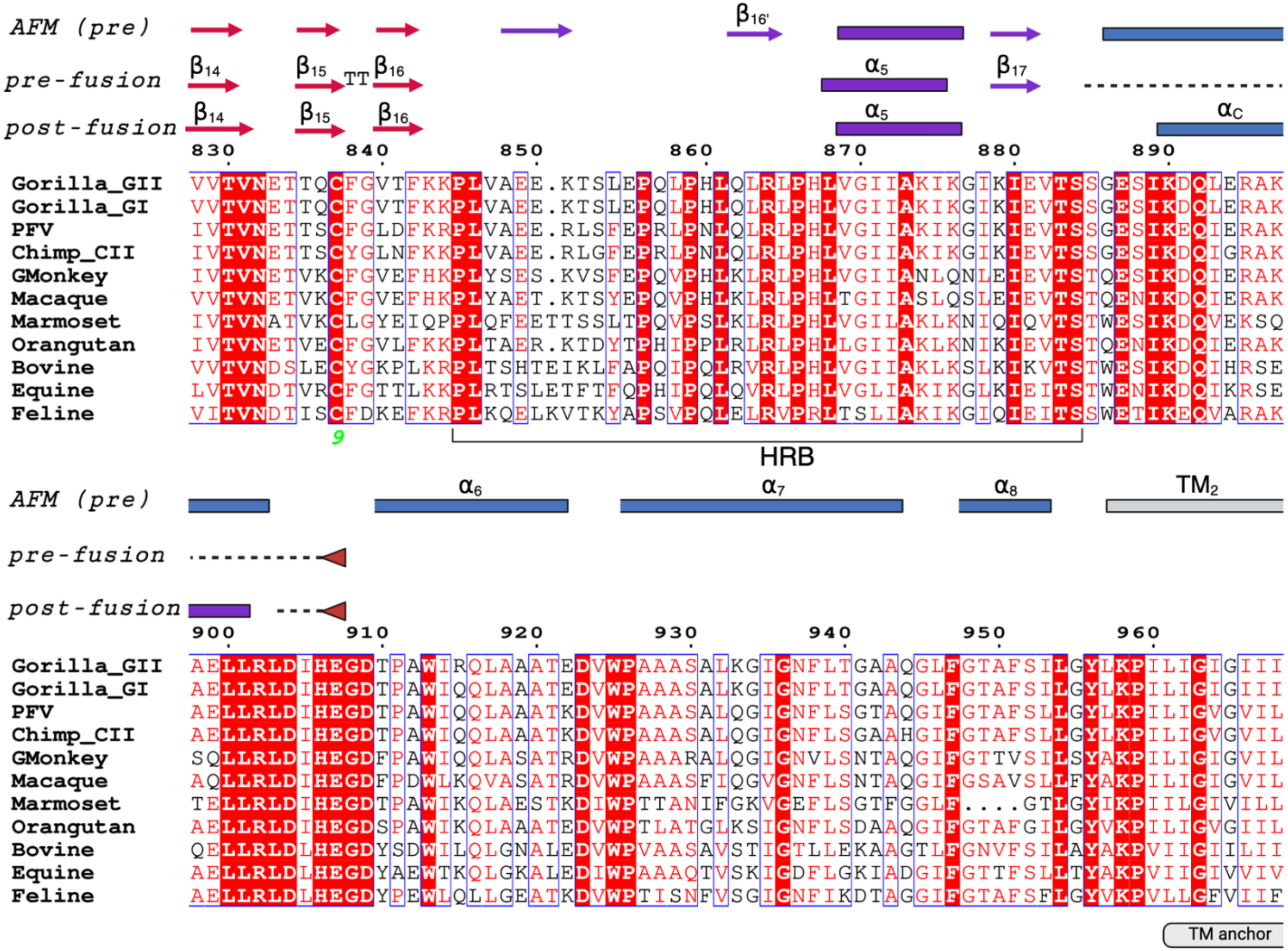
AFM (pre)
pre-fusion
post-fusion
β16'
α5
β17
αC
β14
β15
TT
β16
830
840
850
860
870
880
890
Gorilla_GII
Gorilla_GI
PFV
Chimp_CII
GMonkey
Macaque
Marmoset
Orangutan
Bovine
Equine
Feline
9
HRB
α6
α7
α8
TM2
900
910
920
930
940
950
960
TM anchor

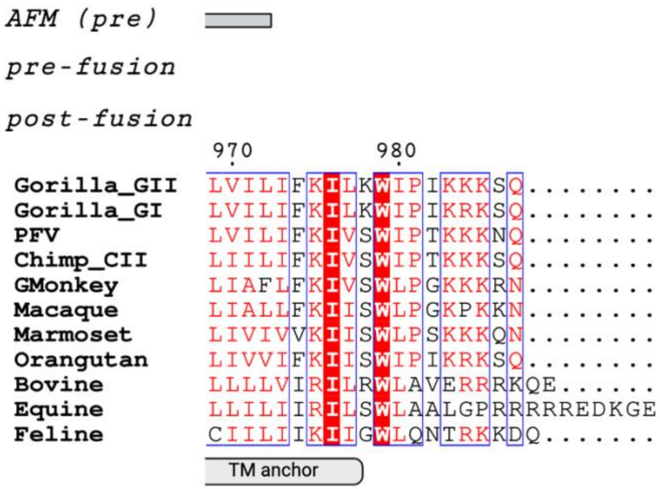

**Figure S6 legend:** Sequences corresponding to 11 FV Env were aligned in Clustal Omega 5 and the alignment was plotted with ESPript https://espript.ibcp.fr 6, with colors that indicate % identity (white letter, red background - identical; red letters, white background identical in >70% sequences; black letters, white background, identical in <70% sequence) [3, 4]. The secondary structure elements corresponding to the Alphafold multimer (AFM) pre-fusion model, the pre-fusion and post-fusion Env ectodomain cryo-EM structures are indicated above the alignment. The color code corresponds to the scheme from Fig. 1C. The $\alpha$-helices and $\beta$-strands present in the AFM model and not in the experimental structures are marked. The terminology for the RBD secondary structure elements was established [5] and the labels were omitted here for clarity. The amino acids constituting the predicted FP (residues 579-596 [6, 7]), HRA (residues 577-651), HRB (residues 845-884) and HRC (residues 186-206) are indicated below the alignment, as well as the membrane spanning segments (TM anchors) we chose based on Phyre 2 [8] prediction of $\alpha$-helical structures.

The Env sequences used in the alignment were obtained from public databases and with following accession numbers: SFVggo_huBAK74 (GII-K74, genotype II gorilla SFV, GenBank: AFX98090.1), SFVggo_huBAD468 (GI-D468, genotype I gorilla SFV; GenBank: AFX98095.1), SFVpsc_huHSRV13 (CI-PFV, known as Prototype Foamy Virus genotype I Eastern chimpanzee SFV; GenBank: AQM52259.1), SFVcpz (genotype I Western chimpanzee; UniProtKB/Swiss-Prot: Q87041.1), SFVcae_LK3 (Genotype II African green monkey SFV; NCBI Reference: YP_001956723.2), SFVmcy_FV21 (genotype I macaque SFV; UniProtKB/Swiss-Prot: P23073.3), SFVcja_FXV (Marmoset FV; GenBank: GU356395.1), SFVppy_bella (Orangutan SFV; GenBank: CAD67563.1), BFVbta_BSV11 (Bovine FV; NCBI Reference: NP_044930.1), EFVeca_1 (Equine FV; GenBank: AAF64415.1), and FFVfca_FUV7 (Feline FV; UniProtKB/Swiss-Prot: O56861.1). Genotypes I and II have been defined for gorilla, chimpanzee, green monkey and macaque FVs [9-11].

#### Figure S7: Inter-protomer disulfide bond connects N-SU and C-SU* of FV Env

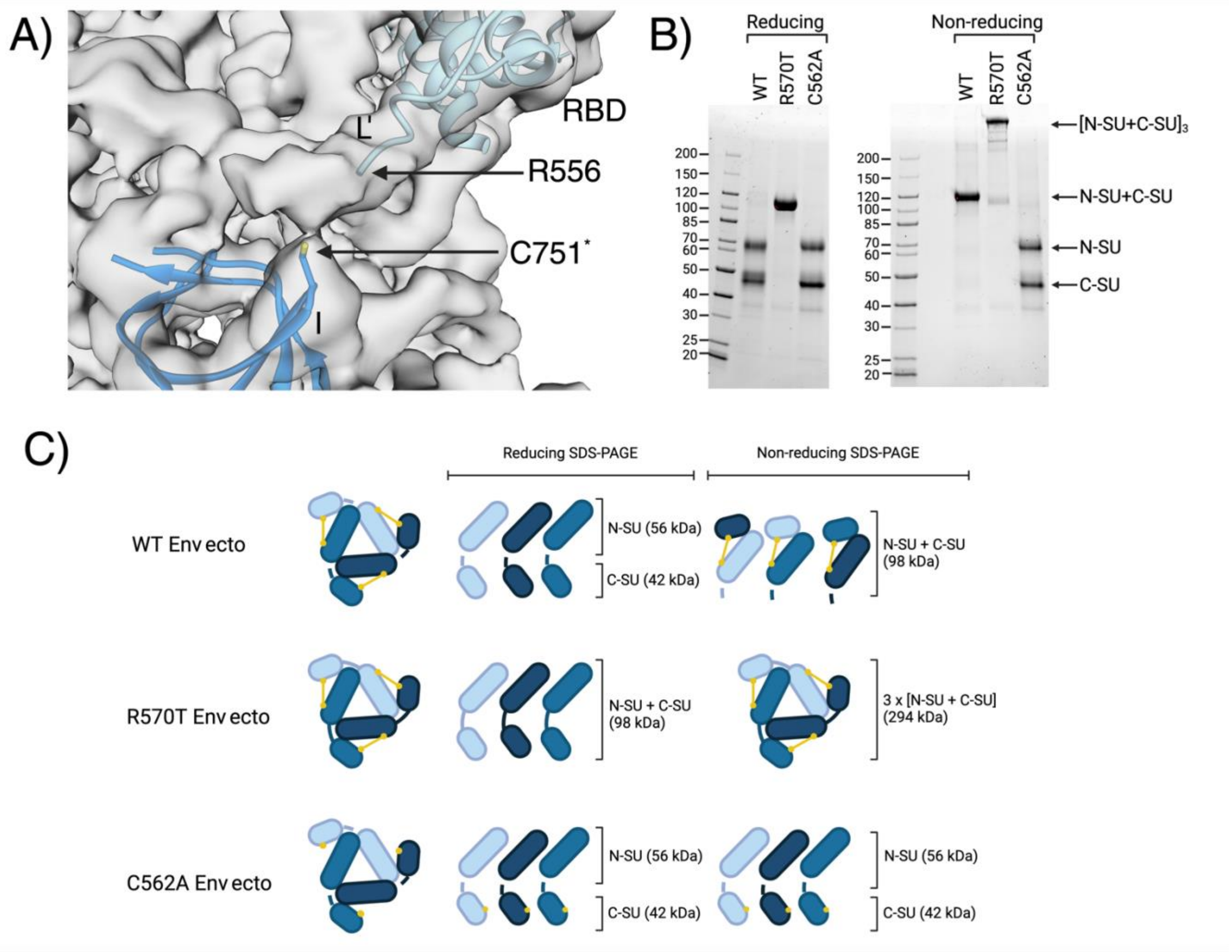

**Inter-protomer disulfide bond connects N-SU and C-SU* of FV Env**

**A)** Unsharpened EM maps calculated for the pre-fusion Env construct V.2 are shown (Fig. S4A), with the RBD belonging to one protomer, and domain I belonging to an adjacent protomer, colored in different shades of blue. The last resolved residue in L', Arg 556 is indicated, as well as Cys 751*, illustrating that the Cys 562 is likely within the reach to form the inter-protomer disulfide bond.

**B)** Coomassie stained SDS PAGE gels of the WT Env ectodomain (fully cleaved at Arg 570), R570T Env ectodomain variant (not cleaved as it lacks the furin site) and C562A variant (fully cleaved because t has the furin site, but is not capable of forming the inter-protomer disulfide bond).

#### Figure S8: AF Multimer model features

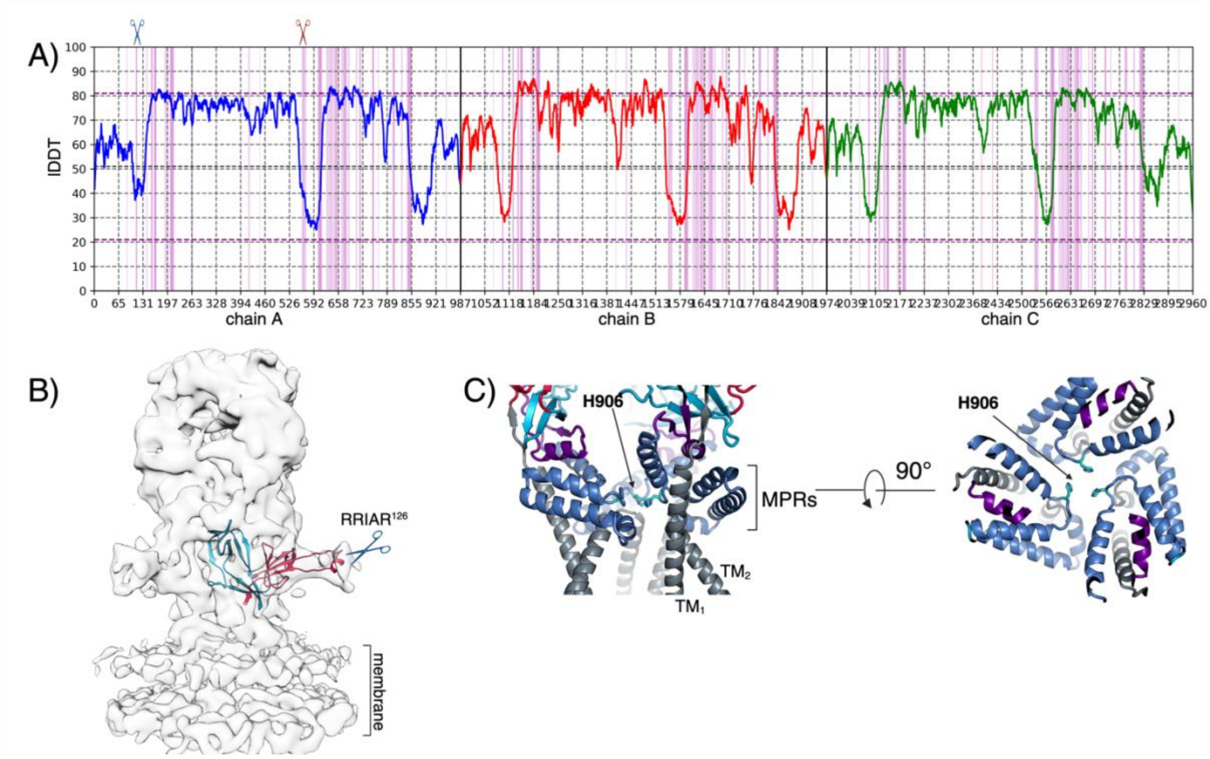

**A)** The lDDT values per residues are plotted for each chain (colored in blue, red and green). The plot was generated by DeepUMQA server [12]. The two cleavage sites are indicated with scissor symbols. The segments around the furin-like cleavage site (residues 105-136), the canonical furin site and FP (residues 555-612), and domain IV (residues 852-909) have lDDT values <50 and should be interpreted with caution.

**B)** The domain I and a part of domain II containing the furin-like cleavage site (scissors symbol) from the FV Env model predicted by Alphafold multimer (Fig. 3) were fitted in the EM maps (EMDB 4013) obtained for the full-length Env displayed on viral vectors [2].

**C)** The side chains of the strictly conserved His 906 are represented with cyan sticks. The structural elements are color according to the scheme shown in Fig. 1C.

#### Figure S9: Viral fusogens constituting the 'Ib' subset of class I fusion proteins

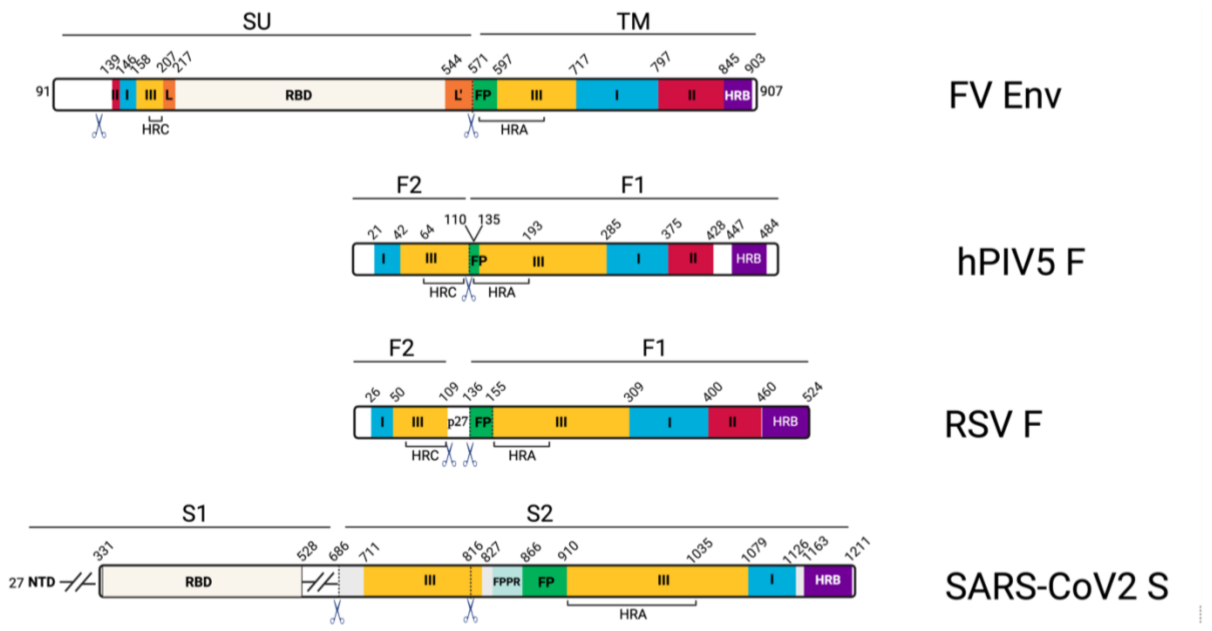

The organization of domains, displayed in a linear manner is shown for the structures displayed in Figure 4. The N-SU and C-SU subunits are labeled with names that had been assigned to each protein in the literature (SU and TM in Env, F2 and F1 in F and S1 and S2 in Spike). The domains are colored according to the scheme in Figure 1C. Furin sites are indicated with the scissors symbol. The S1 subunit of coronavirus Spike is shown in a simplified manner and does not include the N-terminal domain (NTD) or the S1 segment downstream of the RBD. FPPR stands for FP proximal region, structure present only in the Spike. Domain boundaries for the F and S proteins are shown as reported [13-15].

#### Table S1: Cryo-EM data collection, refinement, and validation statistics

| ***Cryo-EM data collection, refinement, and validation statistics*** | | |
|---|---|---|
| | Simian Foamy Virus Env ectodomain **post-fusion** (PDB: 8RM1) | Simian Foamy Virus Env ectodomain **pre-fusion** (PDB: 8RM0) |
| **Data collection and processing** | | |
| Magnification | 105,000x | 150,000x |
| Voltage (kV) | 300 | 200 |
| Microscope | Titan Krios | Glacios |
| Camera | Gatan K3 | Falcon 4i |
| Electron exposure ($e^-$/Å$^2$) | 50 | 40 |
| Defocus range (μm) | -1.0 to -3.0 | -1.0 to -2.75 |
| Pixel size (Å) | 0.85 | 0.95 |
| Symmetry imposed | C3 | C3 |
| Final particle images (number) | 98000 | 15000 |
| Map resolution (Å) | 3.1 | 3.8 |
| FSC threshold | 0.143 | 0.143 |
| **Refinement** | | |
| Initial model | PDB: 8AIC | Post-fusion model + Alphafold model |
| Model resolution (Å) | 2.8 | - |
| Model composition | | |
| Non-hydrogen atoms (number) | 17937 | 16311 |
| Protein residues (number) | 2187 | 2088 |
| Ligands (number) | BMA: 3 | BMA: 3 |
| | NAG: 15 | NAG: 39 |
| | MAN: 0 | MAN: 6 |
| r.m.s. deviations | | |
| Bond lengths (Å) | 0.003 | 0.003 |
| Bond angles (°) | 0.671 | 0.656 |
| Validation | | |
| Molprobity score | 1.63 | 1.50 |
| Clash score | 5.46 | 5.19 |
| Rotamer outliers (%) | 0.00 | 0.19 |
| Ramachandran plot | | |
| Favored (%) | 95.1 | 96.6 |
| Allowed (%) | 4.9 | 3.4 |
| Outliers (%) | 0.0 | 0.0 |

#### Table S2: Superpositions of Env domains from the pre-fusion cryo-EM structure and the AFM model

***Superpositions of Env domains from the pre-fusion cryo-EM structure and the predicted AFM model***

| | Residues range | Total $C_\alpha$ | Aligned $C_\alpha$ | RMSD (Å) |
|---|---|---|---|---|
| dI | 145-161, 713-796 | 82 | 64 | 0.96 |
| dII | 139-144, 797-844 | 48 | 43 | 0.89 |
| dIII | 162-206, 597-712 | 123 | 100 | 1.08 |
| RBD | 217-543 | 320 | 297 | 1.36 |

The superpositions for $C_\alpha$ atoms were carried out using *align* command in Pymol [16]. The domain boundaries are indicated in the 'residue range' column. The values for total and aligned $C_\alpha$ atoms are different in some cases due to different number of residues that were built in the cryo-EM structure and the AFM model. The RMSD stands for the root-mean-square deviation of $C_\alpha$ positions.

#### Table S3: Movement of Env domains during transition from pre- to post-fusion conformation

***Movement# of Env domains during transition from pre- to post-fusion conformation***

| | Residues range | Rotation (°) | Translation vector (Å) | Centre of mass movement (Å) |
|---|---|---|---|---|
| dI | 145-160, 714-796 | 13.06 | 0.41 | 6.20 |
| dII | 139-144, 797-844 | 31.57 | 3.56 | 8.62 |
| dIII | 161-206, 631-713 | 1.77 | 0.01 | 0.42 |
| RBD | 218-543 | 17.35 | 0.51 | 30.61 |

#The movements were measured relative to the helical core and core $\beta$ sheet of domain III, which were fixed for superposition of the pre- and post-fusion structures. The superposition was carried out with the *align* function in Pymol, and the movement parameters were calculated using *draw_axis* command in Pymol [16].

#### Table S4: Superpositions of Env domains from the pre- and post-fusion conformations

| ***Superpositions of Env domains from the pre- and post-fusion conformations*** | | | | |
|---|---|---|---|---|
| | Residues range | Total $C_{\alpha}$ | Aligned $C_{\alpha}$ | RMSD (Å) |
| dI | 145-160, 714-796 | 97 | 68 | 0.78 |
| dII | 139-144, 797-844 | 54 | 54 | 0.98 |
| dIII | 161-206, 631-713 | 129 | 123 | 1.34 |
| RBD | 218-543 | 327 | 275 | 0.65 |

The superpositions for $C_{\alpha}$ atoms were carried out using *align* command in Pymol [16]. The domain boundaries are indicated in the 'residue range' column. The values for total and aligned $C_{\alpha}$ atoms are different in some cases due to different number of residues that were built in the pre- and post-fusion structures. The RMSD stands for the root-mean-square deviation of $C_{\alpha}$ positions.
